# A Dimeric Rocaglate Promotes Multivalent eIF4A-RNA Assembly

**DOI:** 10.64898/2026.03.09.710667

**Authors:** Jie Liu, Megan K. Moore, Kevin Lou, Douglas R. Wassarman, Abolfazl Arab, Samuel Ojeda, Barbara Karakyriakou, Ann-Sophie Koglin, Christopher J. Ott, Luke A. Gilbert, Kevan M. Shokat

## Abstract

Ligand dimerization represents a powerful strategy to enhance avidity, potency, and selectivity. Leveraging the natural-product molecular glue Rocaglamide (RocA), we identified BisRoc, a dimeric rocaglate ligand that potently and durably suppresses translation and exhibits greater specificity across a cancer cell line panel than the monomeric RocA. CRISPRi screening revealed that BisRoc activity is influenced by cellular context, including IFITM-mediated uptake, ABC-type efflux transporters, and the translation initiation factor eIF4A2. Mechanistic studies showed that the paralogs eIF4A1 and eIF4A2 are differentially sensitive to BisRoc-induced dimerization. Owing to the presence of multiple binding sites on RNAs, BisRoc-bridged eIF4A-RNA motifs assemble into higher-order complexes that promote stress-granule formation more efficiently than monomeric RocA. Given the widespread multivalency of RNA-RBP interactions, this ligand dimerization strategy may be extended to modulate the higher-order assembly of other RNA-binding proteins.

**GRAPHICAL ABSTRACT:** 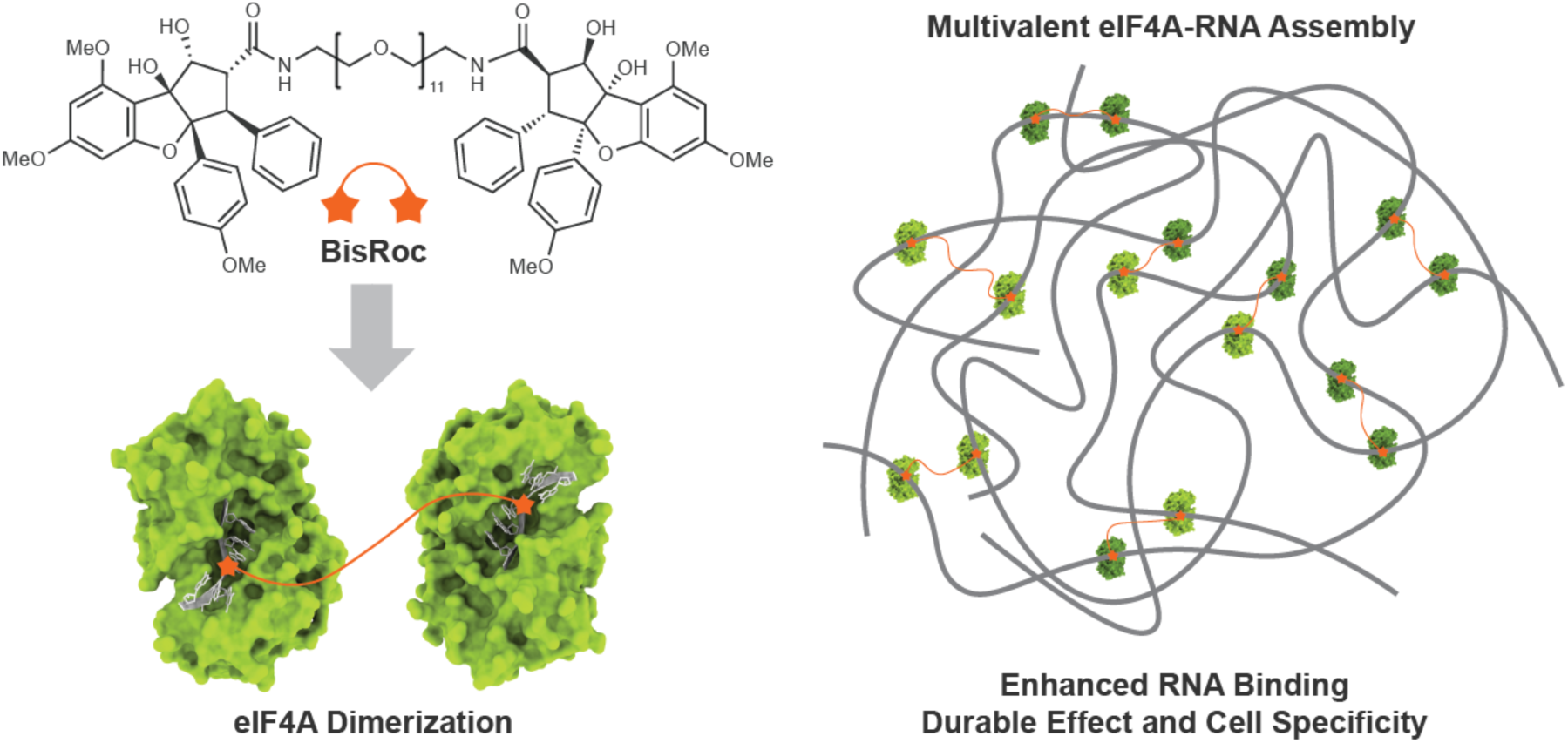

## INTRODUCTION

Eukaryotic translation initiation factor 4A1 (eIF4A1) is an ATP-dependent DEAD-box helicase which plays a critical role in 5’ mRNA cap-dependent protein synthesis.^1–3^ Multiple eIF4A1 molecules are thought to be recruited to 5’-untranslated regions (5’-UTRs) to unwind secondary structures and facilitate 43S preinitiation complex (PIC) scanning and translation initiation.^4–6^ Oncogenic transcripts such as MYC, Cyclin D1, and MCL-1 possess highly structured 5’-UTRs, rendering their translation sensitive to eIF4A1 availability.^7,8^ Elevated eIF4A1 expression has been observed in MYCN-amplified neuroblastoma, which underscores the reliance of certain tumors on this helicase.^9^ This dependency on eIF4A1 has motivated efforts to identify small molecules capable of modulating its activity.^10–12^

Rocaglates are a class of eIF4A1-targeting natural products including rocaglamide (RocA) that exhibits low-nanomolar potency across diverse cancer cell lines.^13–15^ Rocaglates act as molecular glues that clamp eIF4A1 onto polypurine (AG-rich) RNA motifs, thereby stalling protein synthesis.^16^ Ribosome profiling of RocA-treated cells together with eIF4A1 iCLIP data revealed multiple eIF4A1 binding sites across both 5’-UTRs and coding regions.^17^ In parallel, cryo-EM structures of human 48S translation initiation complex showed the presence of two eIF4A1 molecules.^18^ These observations raise the question of whether increasing RocA valency could further enhance the potency or selectivity of eIF4A inhibition.

Ligand dimerization is a well-established strategy for increasing avidity and modulating protein activity.^19^ This approach has been successfully applied across diverse target classes, including cell-surface proteins,^20^ E3 ligases,^21^ inhibitor-of-apoptosis proteins (IAPs),^22^ transcription factors,^23^ and bromodomain and extra-terminal family proteins (BETs).^24,25^ For example, Bradner and coworkers showed that dimeric BET inhibitor MT1 is >100-fold more potent than the monovalent inhibitor JQ1.^24^ More recently, London, Levy, and colleagues demonstrated that dimeric ligands can further drive the polymerization of homomeric targets.^26^ Owing to the symmetric nature of these targets, dimeric ligands can amplify bivalency to multivalency. Considering the reported multivalent eIF4A1-RNA interactions,^6,17,18^ we hypothesized that rocaglate dimerization could promote higher-order eIF4A1-RNA assemblies, which might display activities distinct from monomeric rocaglates.

In this work, we characterize the cellular activity of BisRoc, a dimeric rocaglate analog we recently reported in the context of the mechanism by which large molecular-weight molecules enter cells.^27^ By inducing eIF4A dimerization on RNA, BisRoc exhibits more sustained activity compared to the monomeric control following drug washout. A genome-wide CRISPRi screen identified the cellular import factor IFITM1 and the drug efflux pump ABCC1 as key gene dependencies consistent with the large molecular weight of the dimeric ligand. Unexpectedly, BisRoc displayed a strong dependency on eIF4A2, in contrast to RocA. Target engagement assays showed that BisRoc retained the canonical rocaglate target preference for eIF4A1 and eIF4A2, while promoting stronger cellular engagement of eIF4A2. In vitro experiments revealed distinct BisRoc-induced dimerization sensitivities of eIF4A2 versus eIF4A1 on RNA, driven by a single amino acid difference at the protein-RNA interface. Cellular assays showed that BisRoc promotes higher-order eIF4A-RNA assemblies. At higher concentrations, BisRoc further enhances condensate formation. Together, these findings reveal that a dimeric rocaglate bridges eIF4A-RNA motifs, driving higher-order assemblies that result in stronger and more sustained cellular activity.

## RESULTS

### BisRoc’s bivalency confers potent and durable translational repression

Inspired by the multivalent interaction between multiple copies of eIF4A1 on a single mRNA, we designed a dimeric rocaglate, BisRoc, in which two rocaglate units were linked by a PEG11 spacer to enhance multivalent binding (Figure 1a).^27^ Previous studies have shown that rocaglates preferentially bind polypurine motifs.^16^ We therefore asked whether this enhanced multivalency could further confer sequence-selective inhibition of mRNA translation.^28^ To address this, we performed a puromycin incorporation assay to assess effects on global protein synthesis (Figure 1b).^29^ Puromycin serves as an indicator of protein synthesis that incorporates into the nascent peptides, leading to premature chain termination. The level of puromycin incorporation provides a quantitative readout of ongoing protein synthesis. Treatment with RocA robustly inhibited global protein translation in this assay. The dimeric ligand BisRoc exhibited inhibitory activity comparable to RocA.

**Figure 1.**
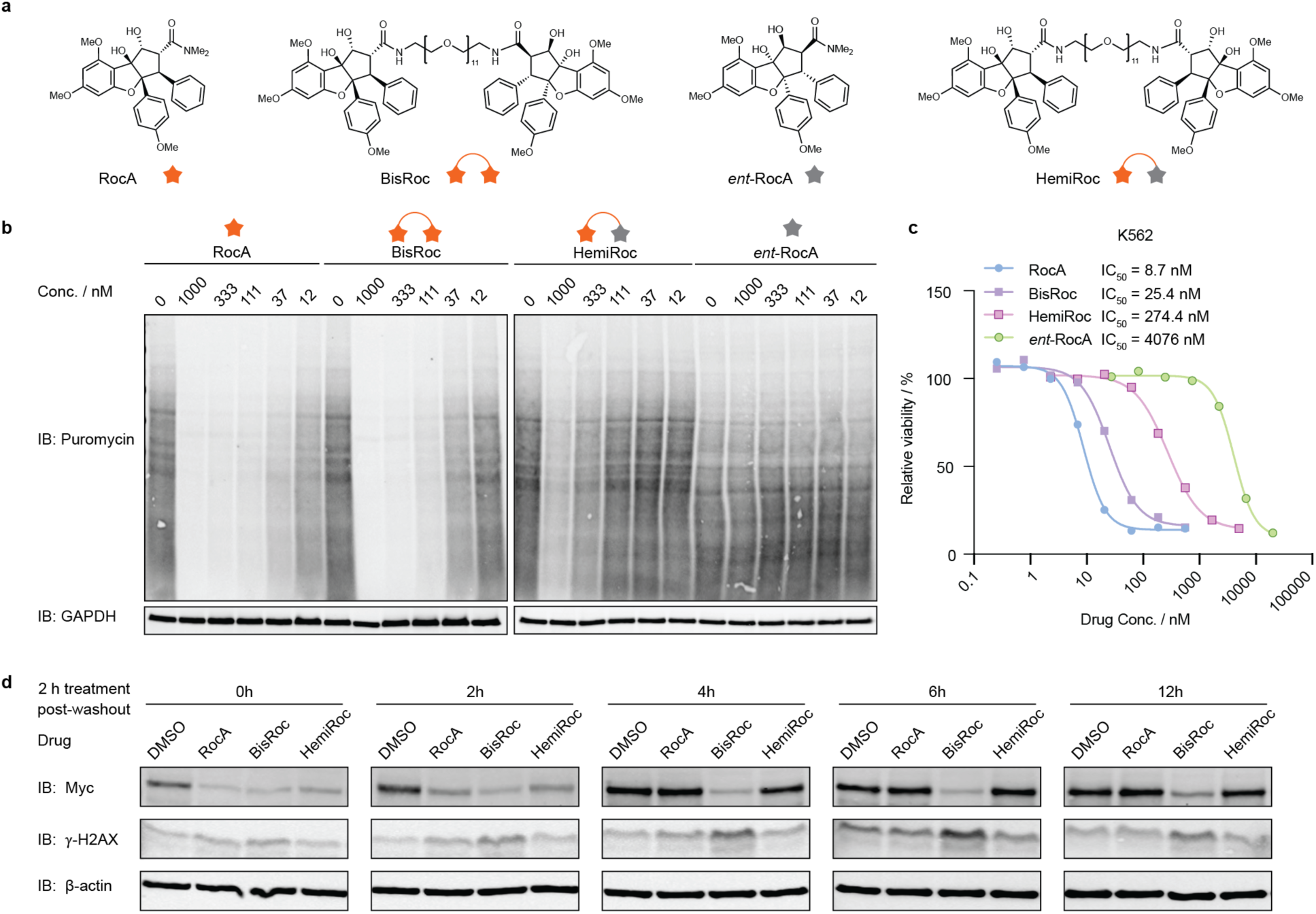
BisRoc exhibits potent and more durable translation suppression than HemiRoc. a. Chemical structures of RocA, BisRoc, *ent*-RocA, and HemiRoc. b. Puromycin incorporation assay comparing RocA, BisRoc, HemiRoc, and *ent*-RocA across a range of concentrations. c. CellTiter-Glo viability assay performed in K562 cells for 72 h. Data are presented as mean for three independent replicates. d. Washout assay (1 µM for drug treatment) on K562 cells assessing the durability of effect on Myc and γ-H2AX.

Western blotting provides only a semiquantitative readout of global translation. To quantitatively compare RocA and BisRoc, we performed translatome proteomics using O-Propargyl-Puromycin (OPP) (Figure S1a).^30^ After OPP incorporation, a biotin tag was introduced by copper-catalyzed click chemistry. The labeled nascent proteome was then enriched, digested, tagged by TMT10 reagents, and analyzed by quantitative mass spectrometry. RocA and BisRoc both exhibited highly correlated effects on nascent protein synthesis (Pearson r = 0.81), indicating largely overlapping sequence selectivity.

Given the similar nascent proteome selectivity of RocA and BisRoc, we next examined whether BisRoc’s bivalent architecture contributes to its cellular potency by generating HemiRoc, in which one rocaglate moiety of BisRoc was replaced by its inactive enantiomer, thereby disrupting bivalency but having the same molecular weight and atomic composition as BisRoc (Figure 1a). The resulting HemiRoc serves as a functionally monomeric control, allowing us to study the specific contribution of the dimeric ligand. *ent*-RocA has been reported as an inactive control for natural RocA,^31^ and this was confirmed in our puromycin incorporation assay, where no inhibition of protein translation was observed (Figure 1b). HemiRoc’s inhibition of global translation was reduced by more than 10-fold relative to BisRoc. We also evaluated the effects of these compounds on cell viability (Figure 1c). RocA and BisRoc strongly inhibited K562 viability, with half-maximum inhibitory concentrations (IC_50_) of 9 and 25 nM, respectively. Consistent with the puromycin incorporation results, HemiRoc exhibited markedly reduced activity, with an 11-fold loss of potency, whereas *ent*-RocA had no significant effect on cell viability at concentrations below 1 µM (Figure 1c). Together, these results support the conclusion that both active rocaglate moieties of BisRoc, which likely engage two target sites, mediate BisRoc’s potent cellular effects.

Because target dimerization can increase avidity and, consequently, target residence time, BisRoc’s activity was evaluated using a cellular washout assay. K562 cells were treated with 1 µM DMSO, RocA, BisRoc, or HemiRoc for 2 h, after which the compounds were removed by replacing the cell medium. Cells were then cultured for the indicated times and pulsed with puromycin for 1 h to assess global translation (Figure S1b). At the starting point, protein translation was strongly suppressed by RocA, BisRoc, and HemiRoc. After 6 h of post-washout incubation, protein translation recovered to the DMSO level in RocA and HemiRoc pretreated cells, however, BisRoc pretreatment resulted in sustained suppression of protein synthesis. This sustained inhibition was still evident even after 12 h of post-washout incubation. Myc downregulation by BisRoc was also sustained for up to 12 h after washout, whereas Myc levels recovered to DMSO control levels within 4 h in RocA- and HemiRoc-pretreated cells (Figure 1d). A recent study by Irish and coworkers demonstrated that rocaglates trigger a robust DNA damage response in leukemia cells, characterized by elevated γH2AX levels.^32^ Consistent with this observation, BisRoc pretreatment in our system led to a prolonged increase in γH2AX that remained elevated for at least 12 hours following compound washout (Figure 1d). Together, these results indicate that although BisRoc and RocA exhibit similar nascent proteome selectivity, BisRoc confers markedly prolonged functional suppression of translation following compound removal. This sustained activity is dependent on its bivalent architecture, as the monovalent control HemiRoc lacks comparable potency and durability.

### BisRoc’s activity is dependent on eIF4A2 and on cellular uptake/efflux pathways

To systematically characterize the cellular mechanisms mediating BisRoc-induced target dimerization, we performed an unbiased chemical-genetic screen to identify differential interactions with monomeric versus dimeric rocaglate molecular glues. CRISPR interference (CRISPRi) is a systematic genome-scale screening method.^33–35^ It not only reports on-target engagement but also reveals critical pathways underlying a drug’s mechanism of action. Using this approach, interferon-induced transmembrane proteins IFITM1-3 were previously identified as key proteins that promote the cellular uptake of large molecules, including BisRoc.^27^ Consequently, IFITMs were expected to emerge as hits in our CRISPRi screen for BisRoc.

The CRISPRi screen was conducted in pre-engineered K562 cells (i.e., K562 CRISPRi cells) that stably express CRISPRi machinery (Figure 2a). A genome-scale sgRNA library was transduced into K562 CRISPRi cells and selected with puromycin, after which the resulting cell pool was treated with DMSO, RocA (10 nM), or BisRoc (5 nM), respectively. During the first 3-day treatment, RocA and BisRoc exhibited comparable inhibition of cell growth (Figure S2a). Beyond Day 3, however, BisRoc consistently and more potently suppressed cell growth relative to RocA. To ensure sufficient cell numbers for sequencing analysis, BisRoc was removed on Day 7, though a sustained inhibition on cell growth was still observed. At the end of the screen, relative sgRNA abundance was quantified by next-generation sequencing (NGS) and gene-level phenotypes were determined using established methods.^34^

**Figure 2.**
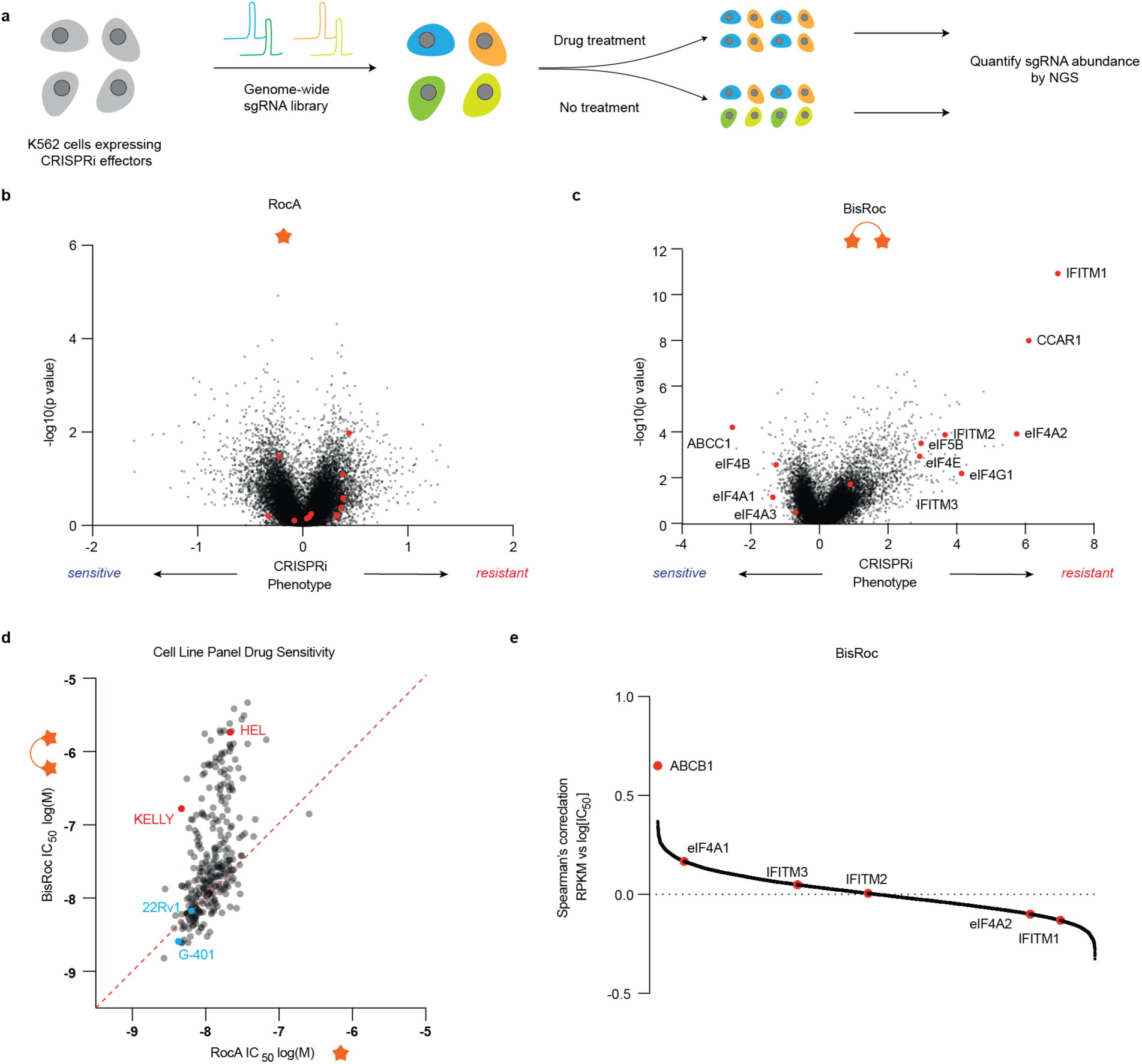
Functional genomics screening reveals distinct chemical-genetic interactions between RocA and BisRoc. a. Workflow of CRISPRi screening. b, c. CRISPRi screening results for RocA and BisRoc. d. Cell line panel screening results across 300 human cancer cell lines. e. Correlation of gene expression and BisRoc’s log[IC_50_] values across the cell line panel.

RocA showed a relatively narrow distribution of gene-level phenotype scores, with most values ranging between -1 and 1 (Figure 2b). In contrast, BisRoc displayed a broader range of phenotypic scores across multiple genes (Figure 2c). As expected, BisRoc’s activity was highly influenced by the cellular uptake/efflux pathways, including IFITM1/2 and the ATP-binding cassette subfamily C member 1 (ABCC1) (Figure 2c). In our previous study, we discovered that IFITMs assist the cellular uptake of linked chemotypes, with IFITM knockdown conferring resistance.^27^ In contrast, ABCC1 functions as an efflux pump that exports molecules from the cytosol. Some proximity-inducing modalities such as PROTACs are reported to be sensitive to ABCC1 expression.^36^ Consistent with this, ABCC1 knockdown enhanced BisRoc sensitivity, suggesting that active cellular export mechanisms similarly regulate intracellular concentrations of linked chemotypes with long linkers (PEG11 in this case).

Notably, multiple translation initiation factors emerged as significant chemical-genetic interactors with BisRoc, including eIF5B, eIF4E, eIF4B, and eIF4G1 (Figure 2c). Rocaglates are reported to target eIF4A1 to impair protein translation.^15,16^ However, knockdown of eIF4A1 did not produce a significant sensitizing or resistant phenotype under either RocA or BisRoc treatment conditions, potentially as a consequence of its essentiality for cell growth. Unexpectedly, its paralog eIF4A2 emerged among the top resistance genes for BisRoc. We further validated this finding by individually testing 3 sgRNAs targeting eIF4A2, all of which exhibited resistance to BisRoc treatment, showing an approximately one-fold increase in IC_50_ upon partial knockdown (Figure S2b,c).

CRISPR screens are a powerful way to study a compound’s mechanism of action, but findings from individual cell lines may not be broadly representative. To complement our CRISPRi screen that was performed in a single cancer cell line, we also conducted a dose-response viability screen across a panel of 300 different cancer cell lines (Figure 2d).^37^ RocA demonstrated consistent cytotoxic potency across the panel, while BisRoc displayed substantial variability. For example, G401 and 22Rv1 cells showed comparable sensitivity to RocA and BisRoc. However, HEL and KELLY cells were more sensitive to RocA (Figure S3a-d).

To examine chemical-genetic interactions, we correlated IC_50_ values with transcript abundance across the cell lines (Figure 2e).^38^ High expression of ABCB1, an efflux pump, emerged as the strongest determinant of BisRoc resistance,^39^ which is analogous to our identification of ABCC1 as a hit in the CRISPRi screen and consistent with functional overlap amongst ABC transporters across different cellular contexts (Figure S3e). In addition, trends involving translation factors were observed but did not reach statistical significance (Figure 2e, S3f-g). Notably, lower eIF4A2 expression was weakly associated with reduced sensitivity to BisRoc treatment, consistent in direction with CRISPRi results, though this correlation was modest (Figure 2e).

### BisRoc binds with eIF4A1 and eIF4A2 in cells

The identification of a particular dependency of BisRoc on the eIF4A2 paralog in the CRISPRi screen attracted our attention as the eIF4A1 and eIF4A2 paralogs share 90% sequence identity. Both belong to the DEAD-box RNA helicases, however, eIF4A1 is expressed at much higher levels and serves as a core component of the eIF4F complex.^3^ Although recent reports have shown that eIF4A2 plays distinct roles in translational repression and stress response,^40–42^ it has generally been considered partially redundant to eIF4A1.^43,44^ Biochemically and functionally, rocaglates are reported to clamp eIF4A1 and eIF4A2 with comparable potency.^41,45^

BisRoc’s distinct dependency on eIF4A2, rather than eIF4A1, in our CRISPRi screen first suggested a potential for a distinct paralog binding preference compared with RocA. To test this hypothesis, we needed to directly assess cellular target engagement for RocA and BisRoc. There have been several strategies to profile rocaglate targets, such as pull down,^46^ covalent labeling,^47^ and thermal shift proteomics.^48,49^ However, these methods are typically performed in lysates or require RNA/AMP-PNP (a non-hydrolyzable ATP analog) additives, making it challenging to assess target engagement in living cells.

To address this, we turned to photoaffinity-based proteomic profiling (ABPP) (Figure 3a).^50^ In the co-crystal structure of eIF4A1 with RocA and polyAG RNA, the amide side chain of RocA is solvent-exposed, providing a suitable site for chemical modification.^51^ We introduced a minimal photoaffinity group containing a terminal alkyne and diazirine at this site,^52^ yielding RocA-PAL (Figure 3b). This probe retained robust activity in K562 cells, as shown by CTG assays and western blot analysis of Myc (Figure S4a,b). Upon UV irradiation, this probe forms covalent bonds with its cellular targets. To assess its photocrosslinking capacity, we performed a dose-dependent labeling experiment in which the RocA-PAL labeled proteome was conjugated with TAMRA-azide (Figure 3c). At a concentration of 1 µM, RocA-PAL produced a relatively clean labeling pattern with a major band at around 55 kDa, which is consistent with the expected molecular weight of eIF4A1-3. Increasing the dose enhanced the intensity of this band, but other protein bands emerged as well. Based on these results, a concentration of 5 µM was selected for subsequent competitive ABPP experiments to ensure sufficient labeling while minimizing off-target enrichment. RocA-PAL binding was competitively displaced during cotreatment with RocA or BisRoc at concentrations ranging from 5 µM to 50 µM, leading to reduced intensity of the band around 55 kDa, while other bands on the gel remained largely unaffected (Figure S4c).

**Figure 3.**
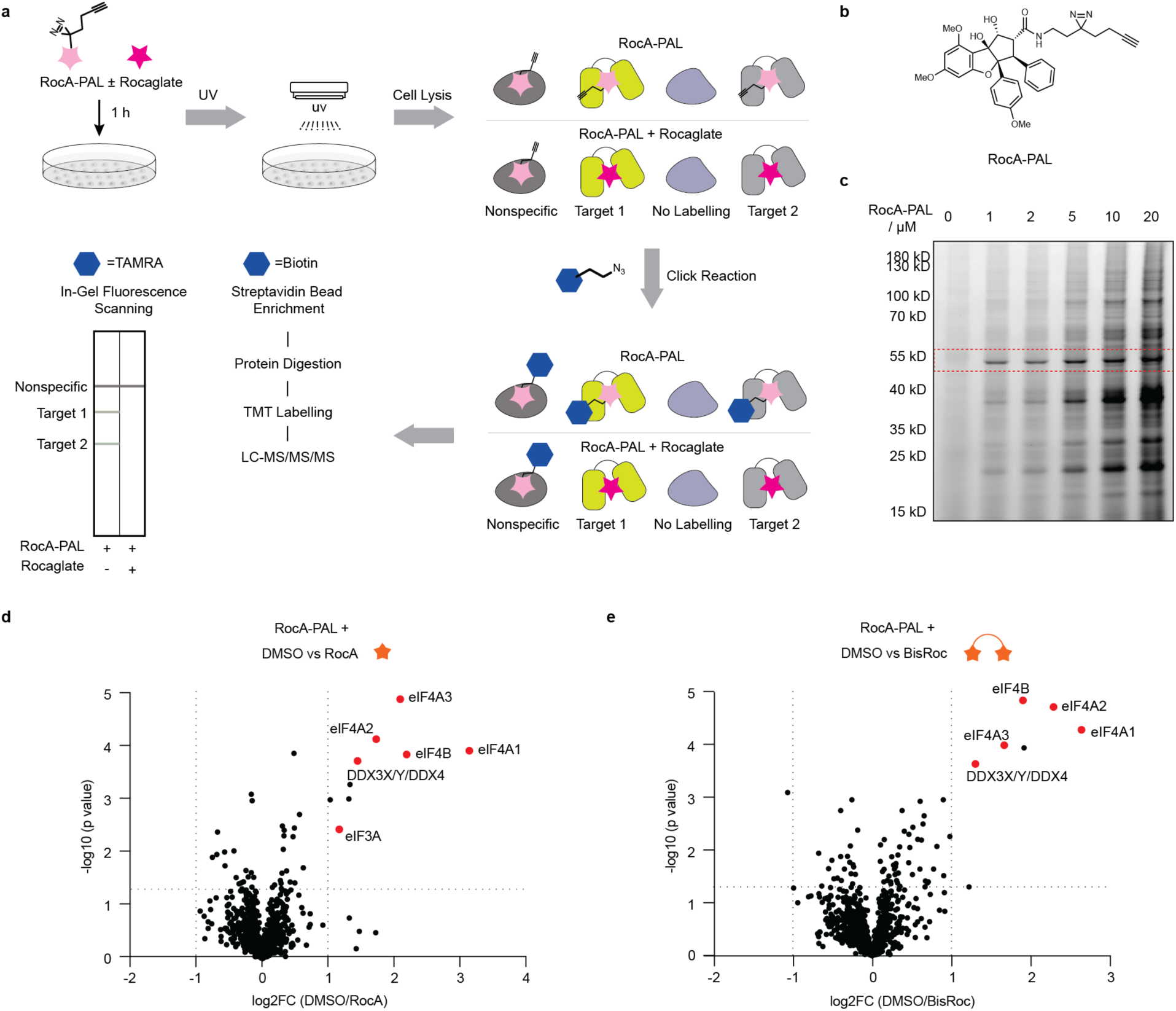
Target engagement evaluation of BisRoc using affinity-based proteomic profiling assay. a. Workflow of affinity-based proteomic profiling (ABPP) experiment. b. Chemical structure of RocA-PAL. c. TAMRA scan of photolabeled proteomes treated with increasing concentrations of RocA-PAL. d, e. Volcano plots from competitive ABPP experiments using RocA or BisRoc as competitors.

To quantify target engagement, photolabeled proteomes co-treated with 50 µM competitors were conjugated with biotin-azide via click chemistry, followed by enrichment, digestion, TMT labeling, and quantitative proteomics. Consistent with previous reports on RocA targets,^46–49^ multiple helicases, including eIF4A1, eIF4A2, eIF4A3, and DDX3X/Y/DDX4, were identified in competitive ABPP experiment using RocA or BisRoc as the competitor (Figure 3d). The translation initiation factor eIF4B also emerged as an interacting partner of rocaglates, likely due to its direct association with eIF4A1 during translation initiation, which may facilitate labeling by RocA-PAL. Although BisRoc did not alter the overall rocaglate target preference as both eIF4A1 and eIF4A2 remained engaged, it produced a modest shift toward eIF4A2 (Figure S4d).

One limitation of competitive ABPP is that it requires relatively high concentrations of RocA and BisRoc. We therefore also employed the cellular thermal shift assay (CETSA), which measures the thermal stability of target proteins upon drug treatment, as an orthogonal method to assess target engagement at lower compound concentrations (Figure S5).^53^ K562 cells were treated with vehicle or 10 µM of RocA or BisRoc and then heated across a temperature gradient. Drug binding is expected to alter the melting temperature of target proteins. Following lysis, soluble fractions were analyzed by western blot against eIF4A1/2. Overall, CETSA results closely mirrored those from ABPP, indicating that BisRoc retained engagement of both eIF4A paralogs, while producing a greater stabilization effect on eIF4A2 than RocA.

### eIF4A1 and eIF4A2 exhibit distinct sensitivities to BisRoc-induced dimerization

When compared with its monomeric control HemiRoc, BisRoc exhibited substantially higher cellular potency (Figure 1c) and slower washout kinetics (Figure 1d), consistent with a dimerization-based mechanism. CRISPRi screening revealed a strong dependency on eIF4A2 rather than eIF4A1. Although cellular target engagement assays indicated that BisRoc retained the canonical rocaglate target preference for both eIF4A1 and eIF4A2, they also suggested enhanced engagement of eIF4A2. Together, these findings motivated us to test whether eIF4A2 is more sensitive to BisRoc-induced dimerization. Although BisRoc’s binding specificity appears relatively unchanged, differential sensitivity to BisRoc-induced dimerization may explain the CRISPRi result and BisRoc’s enhanced activity relative to HemiRoc.

We used a band-shift assay in native gels to evaluate the dimerization efficiency of eIF4A1 and eIF4A2 using poly[AG]_N_ RNA probes of varying lengths, 5’-labeled with 6-carboxyfluorescein (FAM). Under native gel electrophoresis conditions, the protein-RNA-drug complexes remain intact, and the FAM signal attached to the RNA serves as a readout of the complex formation. In native gels, protein migration is primarily determined by size and net charge. When a protein binds negatively charged RNA, the resulting complex carries a higher net negative charge, which can cause it to migrate faster than the apo-protein despite its larger size. Furthermore, higher-order protein-RNA assemblies (containing either additional protein and/or additional RNA components) are up-shifted compared to monomeric complexes (i.e., a single protein bound to a single RNA). This approach allows us to directly compare the ability of BisRoc versus monomeric controls to promote multivalent eIF4A complexes on RNA substrates.

Incubating eIF4A2 and 5’FAM-[AG]_5_ with DMSO, RocA or HemiRoc produced a single band corresponding to a monomeric complex with 5’FAM-[AG]_5_ (Figure 4a). Upon BisRoc treatment, a slower-migrating band appeared, which is consistent with our dimerization hypothesis. With the longer 5’FAM-[AG]_8_ or 5’FAM-[AG]_20_ RNA, BisRoc yielded a prominent slow-migrating band, whereas RocA and HemiRoc produced this band at lower abundance. Because longer RNAs provide multiple eIF4A binding sites, monomeric ligands can also support the formation of larger complexes, but these occur at substantially lower efficiency compared with BisRoc.

**Figure 4.**
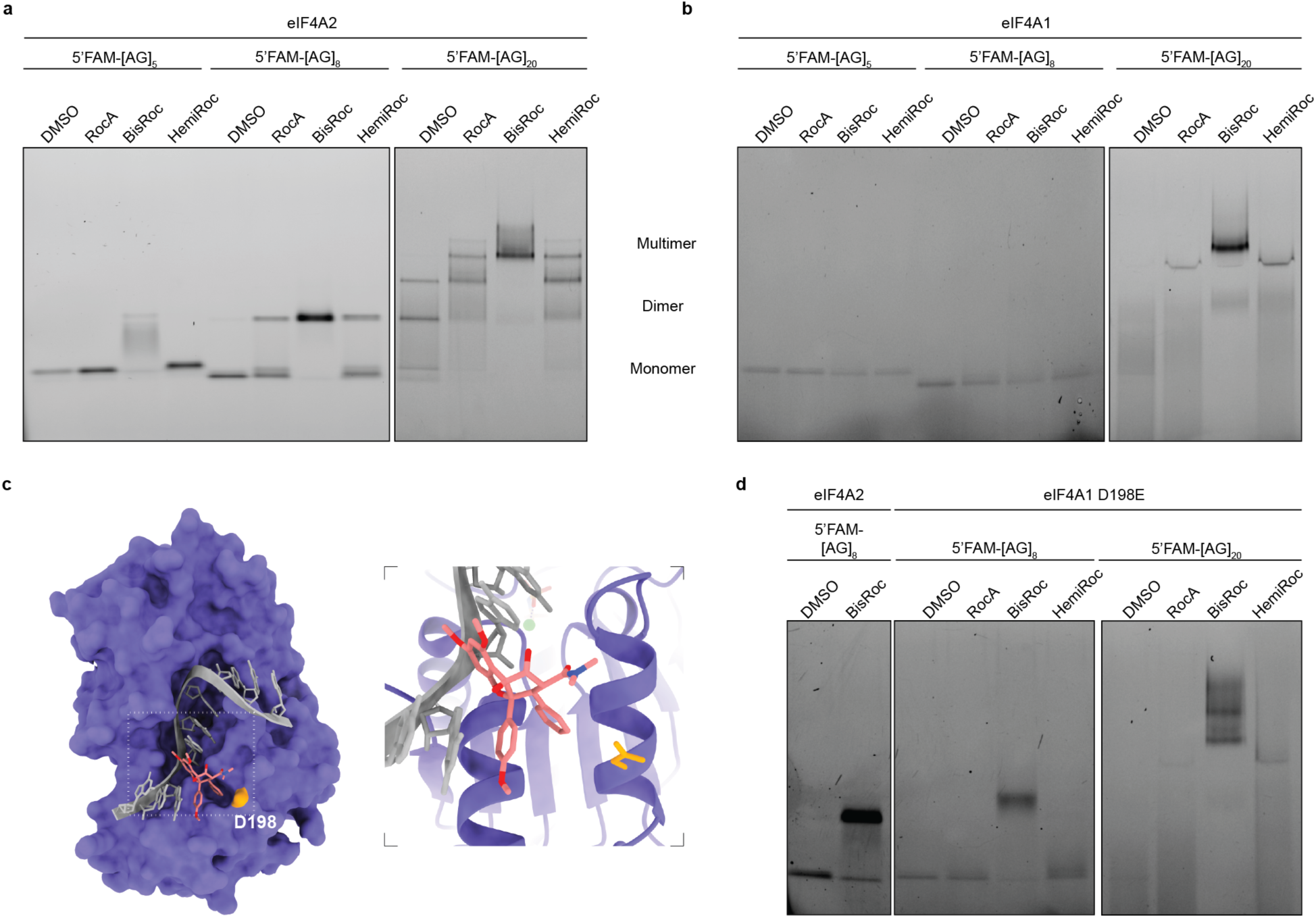
eIF4A2 is more sensitive than eIF4A1 to BisRoc-induced dimerization. a,b. FAM scan of native gel band-shift assay of eIF4A2 and eIF4A1 incubated with poly AG RNAs of varying lengths in the presence of different compound treatments for 30 min. 5’FAM-[AG]_5_ represents the 5’FAM labeled RNA sequence AGAGAGAGAG. Data are representative of two independent experiments. c. Co-crystal structures of eIF4A1-[AG]_5_-RocA (PDB: 5ZC9). Residue D198 was highlighted in orange. d. FAM scan of native gel band shift assays for eIF4A1 D198E. The leftmost two lanes of the eIF4A2 condition were used as references for monomer and dimer bands. Data are representative of two independent experiments.

Since eIF4A1 and eIF4A2 possess nearly identical molecular weights, the bands corresponding to monomeric eIF4A2-[AG]_8_ (DMSO condition) and dimeric eIF4A2-[AG]_8_-BisRoc complexes (BisRoc condition) were used as references to calibrate complex sizes under other conditions. Incubation of eIF4A1 with [AG]_5_ or [AG]_8_ in the presence of compound yielded primarily a single monomeric complex (Figure 4b). In contrast, when the longer [AG]_20_ RNA was used, BisRoc induced the formation of a much larger complex.

The native gel band-shift assay results are consistent with solution-based fluorescence polarization assay results using 5’FAM-poly[AG]_N_ RNA as a probe (Figure S6). When incubated with RNA probe and eIF4A2, BisRoc induced a striking increase in mP values across all different poly[AG]_N_ regardless of length (Figure S6a-c). In contrast, when eIF4A1 was used instead, BisRoc and RocA produced nearly identical polarization signals with [AG]_5_ and [AG]_8_ (Figure S6d,e). When the RNA length was increased to [AG]_20_, we started to observe enhanced mP values upon BisRoc treatment, which suggests higher-order complex formation under such conditions (Figure S6f).

The observed different dimerization sensitivities of eIF4A1 and 4A2 despite their extremely high sequence similarity suggested that one or more non-conserved residues may drive this differential performance. Structural alignment of these two proteins revealed only one amino acid difference at the rocaglate-protein interface: D198 in eIF4A1 versus E199 in eIF4A2 (Figure 4c, S6g). To test the role of this residue, we generated an eIF4A1 D198E mutant and tested it using the same fluorescence polarization assay (Figure S6h) and native gel band-shift assay (Figure 4d). Substitution of Asp198 with Glu resulted in increased mP values and the appearance of a distinct dimer band in the presence of eIF4A1 D198E-[AG]_8_-BisRoc, which phenocopied that by eIF4A2-[AG]_8_-BisRoc condition.

### BisRoc induces an RNA-mediated eIF4A1-eIF4A2 complex in cells, resulting in stress granule

Since the rocaglate-eIF4A interaction is RNA-dependent, the tethering of two eIF4As by BisRoc would bring two RNA motifs together as well (Figure 5a). Given that multiple binding sites exist within a single transcript as well as across transcripts, BisRoc could create an eIF4A1/2-RNA network. To assess this mechanism, we employed a co-immunoprecipitation (CoIP) assay to first determine whether BisRoc induces the formation of such a complex.

**Figure 5.**
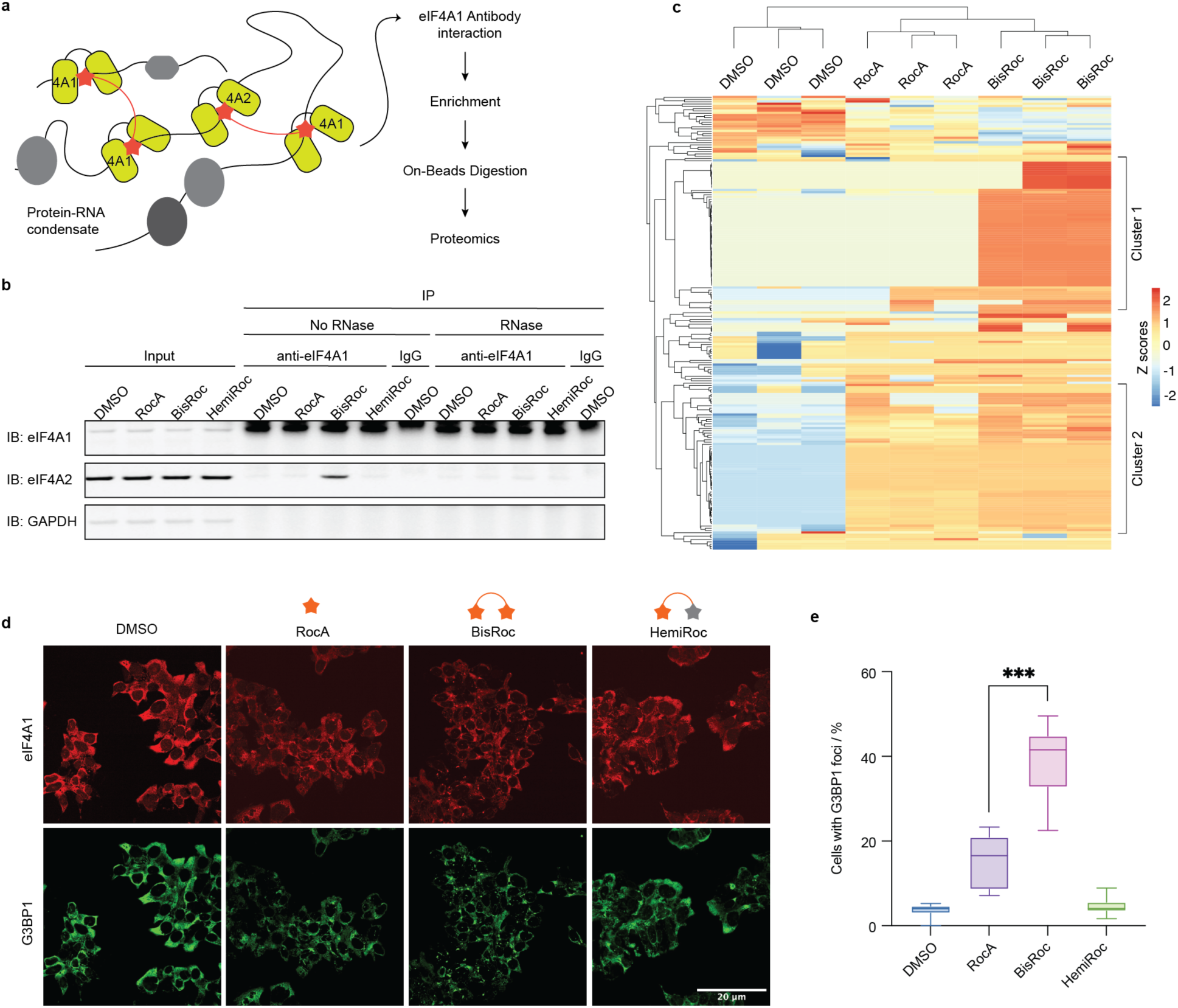
BisRoc promotes higher-order eIF4A-RNA assembly involving eIF4A1 and eIF4A2. a. Workflow of co-immunoprecipitation (CoIP) experiments using anti-eIF4A1 as the bait. b. CoIP results for different drug treatments at 2 µM concentration. Data are representative of two independent experiments. c. CoIP-MS heatmap showing three replicates for each condition. Two clusters were identified: proteins commonly enriched by RocA and BisRoc treatment; proteins uniquely enriched by BisRoc treatment. d. Immunofluorescence microscopy of cells treated with indicated drugs at 1 µM for 24 h. e. Quantification of cells with G3BP1-positive foci. Data are presented as mean ± s.d. (n = 3).

When the anti-eIF4A1 antibody was used as bait, eIF4A2 was co-precipitated under 2 µM BisRoc treatment, whereas no eIF4A2 enrichment was detected under vehicle-, RocA-, or HemiRoc-treated conditions (Figure 5b). This co-immunoprecipitation was RNA-dependent, as eIF4A2 was absent from the elutes upon RNase treatment. Similarly, the anti-eIF4A2 antibody also co-precipitated eIF4A1 under BisRoc treatment (Figure S7a). To investigate the dose-dependency of this phenotype, we performed CoIP experiments across a concentration range from 0.2 nM to 2 µM and found that BisRoc concentrations as low as 20 nM were sufficient to induce co-precipitation of eIF4A1 and eIF4A2 (Figure S7b). Together, these data support the conclusion that BisRoc induces RNA-mediated eIF4A1-eIF4A2 assembly in cells.

To define the composition of the BisRoc-induced eIF4A-RNA network, we performed a CoIP-MS experiment, in which the co-precipitated proteomes were digested and analyzed by mass spectrometry (Figure 5c). Unlike the CoIP/WB results, mass spectrometry did not capture the specific enrichment of eIF4A2 by BisRoc (Figure S7c). Upon closer inspection, we found that some peptides were misassigned by MaxQuant as eIF4A2-unique.^54^ Because eIF4A1 and eIF4A2 share extremely high sequence similarity, with only a single isomeric difference (eIF4A1 I279 vs eIF4A2 L280) within positions 250-300 (Figure S6g), tryptic digestion of this region generates peptides of identical molecular weight. After excluding the four ambiguous I/L-containing peptides, we observed that eIF4A2 enrichment was significantly higher under BisRoc treatment compared with vehicle or RocA (Figure S7d).

We next performed hierarchical clustering analysis of the CoIP proteomics data and identified two major protein clusters. Cluster 1 comprised proteins uniquely enriched by BisRoc and absent from both vehicle- and RocA-treated samples, whereas cluster 2 contained proteins enriched by both RocA and BisRoc relative to vehicle (Figure 5c). STRING cellular component GO analysis revealed that cluster 1 proteins are strongly enriched for cytoplasmic stress-granule and ribonucleoprotein-granule components, whereas cluster 2 is enriched primarily for more generic RNP complexes (Figure S7e, S7f).^55^ These data suggest that BisRoc promotes the accumulation of stress granule–associated ribonucleoproteins within the eIF4A-RNA network.

To validate this stress-granule association, we performed immunofluorescence microscopy to examine representative stress granule (SG) proteins, including G3BP1 and TIAR (Figure 5d).^56^ BisRoc induced strong stress granule formation in approximately 40% of cells at 1 µM (Figure 5e), with G3BP1, TIAR, and eIF4A2 foci colocalizing with eIF4A1 (Figure S8a). In contrast, the monomeric control HemiRoc produced no significant signal, while RocA induced only weak SG formation at the same concentrations. When the concentration was increased from 1 µM to 20 µM, BisRoc displayed stronger SG-inducing activity (Figure S8b). Although RocA possesses comparable inhibitory effects on global protein translation as BisRoc (Figure 1b), its SG-inducing ability is significantly lower than that of BisRoc, indicating that enhanced SG induction by BisRoc cannot be explained simply by stronger translational repression, and instead is associated with its ability to promote higher-order eIF4A-RNA assemblies.^57^

## DISCUSSION

Direct inhibition of oncogenic mRNA translation is a promising therapeutic strategy when selective suppression of a defined subset of transcripts is possible.^58^ Rocaglates enhance the affinity of eIF4A for polypurine sequences, thereby stalling protein translation by preventing 43S preinitiation complex (PIC) scanning. Although recent advances have improved selectivity for RNA motifs and target helicases, strategies to enhance transcript-level and cell-type selectivity remain limited.^28,49,59^

Here, we identified a dimeric rocaglate, BisRoc,^27^ which exhibits markedly greater cell specificity than RocA across a cancer cell line panel. Unbiased CRISPRi screening further revealed that BisRoc’s activity is strongly dependent on cellular uptake and efflux pathways. This dependency has also been observed in multiple proximity-inducing modalities.^27,36,39^ Large molecular-weight chemotypes often exhibit limited passive permeability, increasing their dependency on cellular uptake and efflux pathways.

Furthermore, the CRISPRi screening unexpectedly revealed a unique dependency of BisRoc on eIF4A2. eIF4A2 is generally considered functionally redundant with eIF4A1. Nevertheless, recent studies have shown that eIF4A2 is closely linked to translational suppression through its interaction with the CCR4-NOT complex.^40–42^ In this work, we show that BisRoc’s dependency on eIF4A2 is associated with the greater propensity of eIF4A2, relative to eIF4A1, to undergo ligand-induced dimerization on RNA. Intriguingly, this sensitivity to dimerization is influenced by a single residue at the eIF4A-RNA-rocaglate interface. The D198E mutation in eIF4A1 increases sensitivity to dimerization relative to the wild type.

Rocaglates are thought to inhibit translation through at least two mechanisms: clamping eIF4A onto polypurine-rich 5’-UTR elements to block 43S PIC scanning, and sequestering eIF4A in non-productive complexes, thereby depleting it from eIF4F.^16,60^ Owing to its dimeric architecture, BisRoc may reinforce both processes by bridging two eIF4A-RNA motifs. Given the intrinsic multivalency of eIF4A-RNA interactions,^6,17,18^ BisRoc-mediated bridging may occur at multiple sites within a transcript or across distinct transcripts, thereby promoting higher-order assemblies. Consistent with this model, 20 nM BisRoc treatment was sufficient to induce co-precipitation of eIF4A1 and eIF4A2 in CoIP assays (Figure S7b), supporting a mechanism distinct from that of monomeric rocaglates. These multivalent eIF4A-RNA assemblies may also recruit additional RNA-binding proteins and help sustain stress granule formation. In agreement with this idea, immunofluorescence analysis showed that BisRoc induced stress granules more strongly than RocA.

This work illustrates how dimeric ligand design can be leveraged to achieve context-dependent cell specificity. Multiple layers of cellular gatekeeping shape BisRoc activity: IFITMs influence uptake of large chemotypes, intracellular multivalent engagement stabilizes target assemblies and prolongs residence time, and ABC-type efflux pumps regulate compound retention. Together, these factors create a selective landscape in which dimerization confers durable and context-dependent effects. More broadly, these findings demonstrate that dimeric molecular glue design can reorganize RNA-protein networks through multivalent engagement, revealing a mechanism distinct from monovalent clamping. Given the widespread multivalency of RNA-RBP interactions, rational ligand dimerization may represent a broadly applicable strategy for selectively modulating RNA-RBP networks with improved cellular selectivity.

## METHODS

### Chemical synthesis and compound characterization

Synthetic procedures and characterization data for all newly synthesized compounds are included in the Supplementary Information.

### Recombinant protein expression and purification

The DNA sequences encoding human eIF4A1 WT, eIF4A1 D198E, and eIF4A2 were synthesized by Genscript. They were then cloned into the pET21a(+) vector with an N-terminal TEV-cleavable 6xHis tag.

BL21(DE3) chemically competent cells (NEB, C2527H) were transformed with the pET21a(+)-His-TEV-eIF4A expression plasmids and spread onto LB agar plates containing 100 µg/mL ampicillin. Plates were incubated overnight at 37 °C, and a single resulting colony was used to inoculate Terrific Broth supplemented with 100 µg/mL ampicillin. The culture was grown at 37 °C with shaking at 200 rpm until reaching an OD₆₀₀ of approximately 0.6. Expression was initiated by adding 0.5 mM IPTG at 18 °C for overnight induction. Cells were collected by centrifugation at 4,500 × g for 15 min, resuspended in lysis buffer (20 mM HEPES pH 7.5, 500 mM NaCl, 10 mM imidazole, 2 mM DTT), and lysed by sonication. After clarification by centrifugation at 19,000 × g for 1 h, the supernatant was incubated with Co-TALON resin (Clontech; 4 mL slurry per liter culture) for 1 h at 4 °C with gentle rotation. The resin was washed three times with lysis buffer (10 mL each), and bound His-tagged eIF4A proteins were eluted using 20 mM HEPES pH 7.5, 500 mM NaCl, 250 mM imidazole, and 2 mM DTT (15 mL total).

His-tag removal was performed by overnight digestion with His-tagged TEV protease (Macrolab, UC Berkeley) during dialysis against 20 mM HEPES pH 7.5, 200 mM NaCl, and 2 mM DTT at 4 °C. The digest was briefly incubated with HisPur Ni-NTA resin (Thermo Fisher; 1 mL slurry per liter culture) for less than 10 min, and the unbound fraction containing cleaved eIF4A protein was collected. This material was concentrated to ∼20 mg/mL using a 10 kDa MWCO centrifugal concentrator (Vivaspin Turbo 15, Sartorius) and further purified by size-exclusion chromatography on a Superdex 200 10/300 GL column (GE Healthcare) equilibrated in SEC buffer (20 mM HEPES pH 7.5, 150 mM NaCl, 1 mM DTT). Peak fractions were pooled, concentrated to ∼5 mg/mL, and stored at −78 °C. Typical yields were 5–10 mg purified eIF4A protein per liter of bacterial culture. The mass of the final protein product was verified by LC-MS.

### Fluorescence polarization assays

Purified eIF4A proteins were serially diluted on 384-well plates (Corning, 4514). Compounds and 5’FAM-labeled RNA probes (Genscript) were then added to yield final reaction conditions of 50 µM compound, 10 nM 5’FAM RNA, 20 mM HEPES (pH 7.5), 150 mM NaCl, 1 mM DTT, 1 mM AMP-PNP, and 1 mM MgCl₂ in a total volume of 20 µL. Reactions were incubated at room temperature for 30 min, after which fluorescence polarization was measured on a Spark multimode plate reader (Tecan) at room temperature. mP values were plotted against eIF4A concentrations and binding curve was fit using GraphPad Prism 10.

### Native gel electrophoretic mobility-shift assays

The eIF4A protein (10 µM) and 5’FAM-[AG] RNA probe (100 nM) were assembled with the compound (50 µM) in reaction buffer (20 mM HEPES pH 7.5, 150 mM NaCl, 1 mM DTT, 1 mM AMP-PNP, and 1 mM MgCl₂). After incubation at room temperature for 30 min, 2× native loading buffer (Invitrogen, LC2673) was added, and the mixture was loaded onto an 8–16% native gel (Invitrogen, XP08160BOX). Electrophoresis was performed in Tri-Glycine native running buffer (Invitrogen, LC2672) at 200 V for 40 min. The gel was first scanned for fluorescein signal, followed by Coomassie Blue staining and a second scan to visualize total protein.

### Cell culture

K562, 22Rv1, HEL, KELLY cells (UCSF Cell Culture Facility) were cultured in RPMI 1640 (Gibco, 11995073) supplemented with 4 mM L-glutamine, 1 mM sodium pyruvate, and 10% heat-inactivated FBS (Axenia Biologix). HEK293T cells (UCSF Cell Culture Facility) were maintained in DMEM (Gibco, 11995073) with the same supplements. The K562 CRISPRi (K562i) line was generated as described previously.^34^ G-401 cells (UCSF Cell Culture Facility) were cultured in McCoy’s 5a Medium Modified and 10% heat-inactivated FBS (Axenia Biologix).

All the cells were grown at 37 °C with 5% CO₂ under standard stationary culture conditions unless noted otherwise. All cell lines tested negative for mycoplasma using the MycoAlert Mycoplasma Detection Kit (Lonza).

### DNA transfections and lentivirus production

HEK293T cells were transfected with sgRNA expression vectors and standard packaging vectors (Cellecta, catalog no. CPCP-K2A) using Lipofectamine 3000 Transfection Reagent (Invitrogen, catalog no. L3000015). Lentiviral supernatant was collected 2 days following transfection, filtered through sterile 0.45 µm polyvinylidene difluoride filters (Millipore), and used immediately or stored at -80 °C.

### CRISPRi screening

K562 CRISPRi cells were grown at 37 °C and 5% CO_2_ in 1 L Nalgene Disposable Erlenmeyer flasks with Vented Closure (Thermo Scientific 4112-1000) in a shaking culture (1300 rpm) in a Multitron Incubator (Infors HT). Cells were cultured in RPMI (Gibco) supplemented with 10% fetal bovine serum (FBS) (Avantor Seradigm), penicillin (100 U/mL, Gibco), streptomycin (100 µg/mL, Gibco), glutamine (29.2 µg/mL, Gibco 10378016), and 0.1% Pluronic F-68 (Gibco, Thermo 24040032). Cells were transduced with the 5-sgRNA/gene human CRISPRi v2 library (hCRISPRi-v2) with 8 µg/mL polybrene. Library virus was titered to maximally transduce cells while maintaining a multiplicity of infection (MOI) less than 1. Transduction efficiency and cell growth and viability was monitored daily throughout the experiment by flow cytometry (Attune II, Thermo Fisher). sgRNA+ cells were selected with doses of 1 µg/mL puromycin at 48 and 72 hrs post-transduction until the population was 80-90% sgRNA positive, then cells were allowed to recover in fresh media for 24 hrs before compound treatment. T0 samples were collected before treatment at approximately 750-fold coverage of the library (750 million cells) and the rest of the cells were split into 4 treatment arms with 2 biological replicates per arm: DMSO, 10 nM RocA, 5 nM BisRoc. Cells were cultured at a minimum of 750- to 1000-fold library coverage and continued to be monitored daily by flow cytometry. Dilutions were made daily as necessary with media supplemented with treatments at the indicated concentration to maintain selective pressure until the end of the experiment on Day 11 of compound treatment. The only exception was the BisRoc treatment arms, in which compound was removed after Day 7 of treatment in order to decrease selective pressure to maintain adequate library coverage. Samples were harvested on Day 11 at a minimum of 1000-fold coverage (or 750-fold coverage for BisRoc arms). NucleoSpin Blood XL kits (Macherey-Nagel, Takara Bio 740950.10) were used to extract genomic DNA (gDNA) from all samples and sgRNA protospacers were amplified directly from this with Ti Taq polymerase (Takara Bio 639209), analyzed on an Agilent Tapestation on a High Sensitivity DNA D1000 ScreenTape (Agilent 5067-5584 and 5067-5585), and then sequenced on an Illumina HiSeq 4000 through the UCSF Center for Advanced Technology (UCSF CAT).

### Cloning of single sgRNA expression vectors

sgRNA protospacers targeting individual genes or a negative control were chosen from the library and cloned into the vector pCRISPRia-v2 (Addgene 84832) as previously described.^34^ Complementary oligonucleotides (Elim Bio) were annealed by mixing in Nuclease-Free Duplex Buffer (Integrated DNA Technologies) and heated at 95 °C for 5 min, then cooled at 22 °C for 1 h. The vector pCRISPRia-v2 was digested by BstXI and BlpI (New England Biolabs) and gel purified, then ligated to the annealed duplex with fresh T4 DNA Ligase (New England Biolabs M0202S). Ligated plasmids were transformed into Stellar Competent Cells (Takara Bio 636763) and grown overnight. Individual colonies were grown overnight in 3 mL liquid cultures, plasmids were purified with the Qiagen Spin Miniprep Kit, and sequenced by standard Sanger sequencing (Elim Bio).

### Stable cell line generation

K562 CRISPRi cells (2 x 10^5^) were seeded in 1 mL media in 24-well plates, grown overnight, and treated with lentivirus containing individual sgRNA vectors and 8 µg/mL polybrene (Sigma, TR-1003-G). Cells were treated with 2 µg/mL puromycin (Invivogen, ANT-PR-801) beginning 48 hours post-transduction and puromycin was maintained for 10 days until populations were pure by flow cytometry (CytoFLEX, Beckman Coulter). Knockdowns were checked by immunoblotting over the course of subsequent experiments.

### Cell viability assays

1000 cells were seeded in 100 µL media in 96 well plates and grown overnight. Cells were treated in triplicate with compounds with 0.5% DMSO. Compounds were added by Tecan D300e digital dispenser. Plates were incubated at 37 °C with 5% CO_2_ for 72 h following treatment. 100 µL of a solution of 1:4 CellTiter-Glo (Promega, G7572) to PBS was added per well and plates were shaken for 10 min to mix before luminescence was read out on a Tecan Spark plate reader.

### Cancer cell line panel screening

High-throughput drug screening was performed as previously described.^37^ 300 cancer cell lines were tested for mycoplasma and grown in medium supplemented with penicillin/streptomycin and 10% FBS in a humidified atmosphere at 37°C with 5% CO_2_. All cell lines were grown in RPMI or DMEM/F12 medium to facilitate high-throughput screening and to minimize potential effects of different media on drug sensitivity. Cells were seeded in 384-well plates at varying density to ensure optimal growth of each cell line over the course of the experiment. Cells were treated with vehicle (DMSO) or compound stocks dissolved in DMSO with a PerkinElmer JANUS workstation the day following seeding. Compounds were given in duplicate at 9 doses with a 3-fold dilution factor. The highest dose of RocA was 1 µM and the highest dose of BisRoc was 3 µM. A Day 0 readout was done for growth-rate normalization. Cell viability was determined by CellTiter-Glo after five days of compound treatment and compound-treated wells were normalized to DMSO-treated wells.

### Correlation analysis between cell viability and gene expression

Dose-dependent inhibition of 300 cancer cell lines by RocA and BisRoc was determined first, where dose-response curves were fitted to a four-parameter log-logistic model using SciPy scipy.optimize.curve_fit to estimate IC_50_ values. Correlations of drug sensitivity were determined using the cellpanelr web-app^38^ (https://dwassarman.shinyapps.io/cellpanelr/) with gene expression and proteomics data obtained from the DepMap (22Q1 release) and plotted in GraphPad Prism 10.

### SDS–PAGE and immunoblotting

K562 cells (2 mL medium, 10^6^/mL) were treated with compounds with a final DMSO concentration of 0.5%. At the end of the treatment period, cells were placed on ice. Unless otherwise indicated, cells were pelleted by centrifugation (500g, 5 min), washed once with 1 mL ice-cold PBS, and pelleted again. Unless otherwise specified, cell pellets were lysed on ice for 10 min in RIPA buffer supplemented with protease and phosphatase inhibitors (cOmplete Mini and PhosSTOP, Roche). Protein concentrations were determined using a BCA assay (Thermo Fisher, 23225) and adjusted to 2 mg/mL, or to the highest achievable concentration, with additional RIPA buffer. Lysates were mixed with 4x SDS loading dye (BioRad, 1610747) and denatured at 95 °C for 5 min for immunoblotting.

Unless otherwise specified, SDS–polyacrylamide gel electrophoresis was performed using 4-12% Bis-Tris gels (Novex, Invitrogen) in MES running buffer at 200 V for 50 min. Proteins were transferred onto nitrocellulose membranes using iBlot 3 Transfer Stacks (Invitrogen, IB33002X3). Membranes were blocked in 5% bovine serum albumin (BSA) in Tris-buffered saline with Tween-20 (TBST) for 1 h at 23 °C. Primary antibodies were diluted in 5% BSA–TBST and incubated with membranes at 4 °C for overnight. Membranes were washed three times with TBST (5 min per wash) and incubated with IRDye-conjugated secondary antibodies (goat anti-rabbit IgG-IRDye 800 or goat anti-mouse IgG-IRDye 680; Li-COR) diluted in 5% BSA–TBST for 1 h at 23 °C. Following three additional TBST washes (5 min each), membranes were imaged using a Bio-Rad ChemiDoc Imager.

### Puromycin incorporation assays

K562 cells (2 mL medium, 10^6^cells/mL) were incubated with the drug for 5 hours, followed by the addition of 5 µg/mL puromycin (InvivoGen). The cells were incubated for an additional 60 min before cell harvest and lysis. The western blot was incubated with anti-puromycin to detect the extent of puromycin incorporation.

### Puromycin incorporation proteomics

Similarly, K562 cells (10 mL medium, 10^6^ cells/mL) were incubated with the DMSO, RocA (100 nM), or BisRoc (100 nM) for 5 h (n=3), followed by the addition of 10 µM O-Propargyl-Puromycin (OPP; BroadPharm, BP-50103). Cells were incubated for an additional 60 min prior to cell harvest and lysis in 0.1% NP40, 100 mM HEPES, 150 mM NaCl pH 7.5 with 1x Halt Protease and Phosphatase Inhibitor Cocktails (Fisher, 78447). Cell lysates were clarified by centrifugation (13,000g, 15 min, 4 °C), and protein concentrations were determined using a BCA assay and normalized to 2 mg/mL.

2 mg cell lysates were subjected to click chemistry by conjugation with 0.1 mM biotin-azide (Sigma, 900912-50MG) in the presence of 1 mM CuSO_4_, 0.1 mM TBTA (Vector, CCT-1061), and 1 mM TCEP (GOLDBIO). The reaction was carried out at room temperature for 90 min, after which ice-cold acetone was added, and samples were kept at -20 °C to allow complete protein precipitation. Proteins were pelleted by centrifugation at 4,300 g for 15 min and washed once with cold methanol.

The resulting protein pellet was resuspended in 250 µL 4% SDS in PBS containing 10 mM DTT to ensure complete solubilization, then diluted to a volume of 5 mL with PBS. 100 µL equilibrated NeutrAvidin beads (Fisher, PI29200) were added to the samples. The samples were rotated end-over-end for 2 h at room temperature. Beads were collected by centrifugation, washed once with 0.2 % SDS in PBS, followed by three washes with PBS.

Beads were transferred to protein LoBind tubes (Fisher, 13-698-794) and incubated with 400 µL of 5 mM DTT in 6 M urea/PBS at 56 °C for 30 min. Proteins were then alkylated with 42 µL of 200 mM IAA (Sigma, I6125) for 30 min in the dark. On-bead digestion was performed using 2 µg trypsin (Promega, VA9000) in 100 µL 100 mM TEAB (Sigma, T7408) at 37 °C overnight. The supernatant was collected, and beads were washed twice with 50 µL 100 mM TEAB. All fractions were combined to yield 200 µL of digested peptide solution.

0.8 mg TMT10plex Isobaric Label Reagents (Fisher, 90110) dissolved in 90 µL acetonitrile was added to the peptide solution and incubated for 1 h at room temperature. The reaction was quenched with 12 µL 5% hydroxylamine (Sigma, 467804), followed by the addition of 8 µL formic acid. The labeled peptides were pooled, dried by SpeedVac, and desalted using C18 pipette tips (Fisher, PI87784). Peptides were eluted with three rounds of 100 µL 80% acetonitrile in water containing 0.1% formic acid and dried under vacuum.

### LC-MS/MS acquisition and data analysis of TMT-labeled samples

TMT10plex samples were reconstituted in 50 µL of 5% ACN and 0.1% FA in water and analyzed on an Orbitrap Eclipse Tribrid mass spectrometer (Thermo Fisher Scientific) coupled to an Ultimate 3000 RSLCnano system. Mobile phase A consisted of 0.1% formic acid in water, and mobile phase B consisted of 95% acetonitrile with 0.1% formic acid. For each injection, 5 µL of sample was loaded. Peptides were separated on an EASY-Spray C18 column (3 µm, 75 µm × 15 cm; Thermo Fisher Scientific, Cat. No. ES800) at a flow rate of 0.3 µL min⁻¹. Samples were loaded at 4% B for 20 min, followed by a gradient of 4–10.5% B over 2 min, 10.5–45% B over 73 min, 45–84% B over 2 min, holding at 84% B for 3 min, ramping from 84% to 4% B over 2 min, and re-equilibrating at 4% B for 8 min.

TMT reporter ion quantification was performed using real-time search (RTS)-MS3 methods. MS1 spectra were acquired at a resolution of 120 K with m/z scan range of 375-1800, RF lens of 30%, a maximum ion injection time of 50 ms, charge states of 2-6, and a 2scan30s dynamic exclusion time. MS2 spectra were acquired via HCD at a stepped collision energy of 33 ± 5%, a resolution of 30 K, m/z scan range of 110-3000 and a maximum ion injection time of 75 ms. In RTS-MS3, real-time database searching was performed using the Comet search engine against the reviewed human Swiss-Prot FASTA database with trypsin specified as the digestion enzyme. Methionine oxidation and probe modifications on tyrosine and lysine were set as variable modifications, while carbamidomethylation of cysteine and TMT modifications on N-terminal amines and lysine residues were specified as static modifications. For MS3 acquisition, synchronous precursor selection (SPS) of the top 10 fragment ions was performed, with a maximum injection time of 105 ms. MS3 spectra were acquired in the Orbitrap at a resolution of 50,000 using HCD with a collision energy of 65%.

Peptides were searched against the SwissProt Homo sapiens reference proteome (2021.06.20) using MaxQuant (v.2.0.3.1, https://www.maxquant.org/) with variable methionine oxidation and protein N-terminal acetylation, and fixed cysteine carbamidomethylation. Quantification was performed with reporter ions in MS3. Proteomic statistical analyses were completed using the R programming language, version 4.5.0. Proteins annotated as reverse hits or contaminants were removed, and zero/negative intensities were treated as missing values. For OPP proteomics, Pearson correlation between RocA- and BisRoc-induced log2 fold changes was computed. For ABPP/MS analysis, log2 fold changes and p values were calculated in R using Student’s t-tests (two-tailed, two-sample, equal variance), and final plots were generated using GraphPad Prism 10.

### Washout assays

K562 cells (15 mL medium, 10^6^ cells/mL) were incubated with compounds at 1 µM for 2 h. Compounds were removed by centrifugation at 500 g for 5 min, and cells were resuspended in an equal volume of fresh medium and incubated at 37 °C for 5 min prior to pelleting. This wash step was repeated twice. Cells were then reconstituted in 15 mL fresh medium and aliquoted into separate wells (2 mL each) and harvested at the indicated time points. Cell pellets were lysed and analyzed by western blot.

To analyze the effects on global proteomics, 5 µg/mL puromycin was added at indicated time points and incubated for an additional 1 h before cell harvest.

### Affinity-based protein profiling (ABPP)

K562 cells (10 mL PBS, 10^6^ cells/mL) were treated with the 5 µM RocA-PAL in the presence of DMSO, RocA (50 µM), or BisRoc (50 µM) at 37 °C for 1 h (n=3). Cells were transferred to 4 °C cold room and irradiated with 365 nm UV light (Analytikjena, UVL-225D) for 10 min. Cells were pelleted by centrifugation at 500 g for 5 min, washed once with PBS, and lysed in 1 mL of 0.1% NP40, 100 mM HEPES, 150 mM NaCl pH 7.5 with 1x Halt Protease and Phosphatase Inhibitor Cocktails (Fisher, 78447).

Subsequent sample preparation, enrichment, digestion, TMT labeling, and desalting were performed as described above for puromycin-incorporation proteomics.

For gel-based fluorescent proteomic profiling, cell lysates (24 µg) were subjected to click chemistry by conjugation with 0.025 mM TAMRA-azide (BroadPharm, BP-40173) in the presence of 1 mM CuSO_4_, 0.1 mM TBTA (Vector, CCT-1061), and 1 mM TCEP (GOLDBIO). The reaction was carried out at room temperature for 90 min, after which SDS loading buffer (BioRad, 1610747) was added and samples were incubated at room temperature for 30 min. The samples were resolved by SDS-PAGE, scanned for TAMRA fluorescence, and subsequently stained with Coomassie blue.

### Cellular thermal shift assays (CETSA)

K562 cells (10 mL medium, 10^6^ cells/mL) were treated with DMSO, RocA (10 µM), or BisRoc (10 µM) for 1 h. Then cells were harvested, washed once with cold PBS, and reconstituted to 0.7 mL ice-cold PBS, and divided into 13 aliquots (50 µL each). Aliquots were heated at indicated temperatures for 3 min and immediately flash-frozen in liquid nitrogen. Cells were lysed by three freeze-thaw cycles (liquid nitrogen freezing followed by thawing at room temperature). Lysates were clarified by centrifugation (13,000 x g, 10 min, 4 °C), and 15 µL of the supernatant was mixed with 5 µL SDS loading buffer, resolved by SDS-PAGE, and analyzed by western blot. Relative signal intensities of eIF4A1 and eIF4A2 were normalized to GAPDH, further normalized to the lowest temperature, plotted against temperatures, and fitted using GraphPad Prism 10.

### Co-immunoprecipitation (CoIP)

K562 cells (10 mL medium, 10^6^ cells/mL) were treated with DMSO, RocA, BisRoc, or HemiRoc at the indicated concentrations for 6 h. Cells were pelleted by centrifugation (500g, 5 min, 4 °C), washed once with ice-cold PBS, and lysed in IP lysis buffer (Thermo Fisher, 87788) supplemented with 6 mM MgCl_2_ and 1× Halt protease and phosphatase inhibitor cocktails (Thermo Fisher, 78447). Lysates were clarified by centrifugation (13,000g, 15 min, 4 °C), and protein concentrations were determined by BCA assay and normalized to 1 mg/mL with IP lysis buffer.

Normalized lysates (400 µg) were incubated overnight at 4 °C with 2.5 µg anti-eIF4A1 (Abcam, ab31217), anti-eIF4A2 (Santa Cruz Biotechnology, SC-137148), or IgG control antibodies (Cell Signaling Technology, 3900S or 5415S). Where indicated, RNase A (10 µg/mL; NEB, T3018L) was added during the incubation. Protein A/G magnetic beads (25 µL bead slurry; Thermo Fisher, 88803) were equilibrated, added to the samples, and incubated at room temperature for 1 h. Beads were collected on a magnetic rack and washed twice with IP lysis buffer and once with water.

For western blot analysis, beads were resuspended in 100 µL 1× SDS loading buffer and boiled at 95 °C for 10 min, after which eluates were collected using a magnetic rack.

For proteomics analysis, cells were treated with compounds at a final concentration of 100 nM. Anti-eIF4A1 antibody or IgG control was used as the immunoprecipitation bait. Following the procedures described above, beads were incubated with 50 µL 6 M urea/PBS at 56 °C for 30 min, diluted to 100 µL with 100 mM TEAB, reduced with 7.5 µL 200 mM TCEP at 56 °C for 60 min, and alkylated with 7.5 µL 375 mM iodoacetamide in 200 mM TEAB at 37 °C for 30 min in the dark. 3 µg trypsin in 100 µL 100 mM TEAB was added to the mixture and incubated at 37 °C overnight. Digestion was quenched by addition of 5 µL TFA. Peptides were collected on magnetic rack. The remaining beads were washed twice with 50 µL 100 mM TEAB. The three fractions were combined and desalted using C18 pipette tips (Thermo Fisher, PI87784), eluted with three rounds of 100 µL 80% acetonitrile in water containing 0.1% formic acid, and dried under vacuum.

### LC-MS/MS acquisition and data analysis of label-free samples

Label-free samples were reconstituted in 50 µL of 5% ACN and 0.1% FA in water and analyzed on an Orbitrap Eclipse Tribrid mass spectrometer (Thermo Fisher Scientific) coupled to an Ultimate 3000 RSLCnano system. Mobile phase A consisted of 0.1% formic acid in water, and mobile phase B consisted of 95% acetonitrile with 0.1% formic acid. For each injection, 5 µL of sample was loaded.

Peptides were separated on an EASY-Spray C18 column (3 µm, 75 µm × 15 cm; Thermo Fisher Scientific, Cat. No. ES800) at a flow rate of 0.3 µL min⁻¹. Samples were loaded at 2% B for 12 min, followed by a gradient of 2–31.6% B over 72 min, 31.6–52.6% B over 2 min, ramping from 52.6% to 2% B over 2 min, and re-equilibrating at 2% B for 7 min.

MS1 spectra were acquired at a resolution of 120 K with m/z scan range of 375-1500, RF lens of 30%, a maximum ion injection time of 50 ms, charge states of 2-7, and a 30s dynamic exclusion time. MS2 spectra were acquired via HCD at a stepped collision energy of 30%, in the orbitrap with an isolation width of 1.6 m/z for the label free sample, a resolution of 30 K, auto m/z scan range and a maximum ion injection time of 100 ms.

Peptides were searched according to the process mentioned above. Label-free quantification (LFQ) intensities were processed in R. Raw LFQ values were log-transformed as log2(LFQ + 1), and per-replicate enrichment scores (R1–R3) were calculated as the difference between target and matched control channels for DMSO, RocA, and BisRoc. Enrichment matrices were z-score scaled across samples on a per-protein basis. Hierarchical clustering was performed on both rows and columns using Euclidean distance and complete linkage, and heatmaps were generated using pheatmap.

For focused visualization, the top 200 proteins were selected based on mean BisRoc enrichment across replicates and clustered as above. The resulting row dendrogram was partitioned into eight clusters, and the cluster 1 and 2 were subjected to cellular-component Gene Ontology analysis using STRING (https://string-db.org/).

### Immunofluorescence microscopy

22Rv1 cells (5,000 cells per well) were seeded into 96-well plates (Cellvis P96-1-N, glass thickness 0.13–0.16 mm) and incubated overnight at 37 °C. Cells were treated with compounds or DMSO as a vehicle control for 24 h. Following treatment, cells were fixed with 3.7% paraformaldehyde in PBS and permeabilized with 0.1% Triton X-100 in PBS. After removal of the permeabilization solution, cells were washed with PBS-T and blocked with 100 µL of 2% BSA in PBS-T for 1 h at room temperature. Cells were then stained with DAPI (Sigma-Aldrich, D9542-10MG) to visualize DNA and either anti-eIF4A1 antibody (MedChemExpress, HY-P80650) or anti-eIF4A2 antibody (Abcam, ab31218), anti-G3BP1 antibody (Cell Signaling Technology, 61559), anti-TIAR antibody (Cell Signaling Technology, 8509) overnight at 4 °C. Images were acquired using a CSU-W1 Spinning Disk/High Speed Widefield microscopy equipped with a Plan Apo λ 40x/0.95.

For automated image analysis, the percentage of G3BP1 foci positive cells was determined using CellProfiler software.^61^ Data are presented as mean ± SD (n = 3).

## Acknowledgements

K.M.S. thanks NIH (U19AI171110 & U54 CA243125) and the Sjöberg Foundation for support. We thank K. Herrington from Center for Advanced Light Microscopy at UCSF for her microscopy assistance and expertise (1S10OD017993-01A1). Sequencing on an Illumina HiSeq 4000 through the UCSF Center for Advanced Technology was supported by UCSF PBBR, RRP IMIA, and NIH 1S10OD028511-01 grants. This work was supported by Arc Institute (L.A.G.). We thank Prof. Jack Taunton for the use of the Orbitrap Eclipse. We thank Dr. Siyi Wang and Prof. Davide Ruggero for proofreading this manuscript.

## Contributions

J.L., M.K.M., K.L., and K.M.S. conceived the study. J.L. and K.M.S. wrote the manuscript. J.L. designed and synthesized the tool compounds, expressed proteins, and performed biochemical assays. K.L. provided BisRoc sample initially. M.K.M., K.L., and L.A.G. conducted CRISPRi screening and data analysis. A.A. performed re-analysis of the CRISPRi data. S.O., B.K., A.K., and C.J.O. conducted cancer cell line panel screening and data analysis. J.L., M.K.M. and D.R.W. performed gene dependency analysis. J.L. performed all other cellular assays, microscopy, proteomics, and data analysis. All authors edited and approved the manuscript.

## Data availability

Uncropped gel images are provided as Source Data files accompanying this paper. All proteomic raw data have been deposited to MassIVE (http://massive.ucsd.edu) with accession no. MSV000101038, as well as in ProteomeXchange (http://www.proteomexchange.org) with accession no. PXD075236. Source data are provided with this paper.

## Code availability

Analysis code for OPP proteomics, ABPP proteomics, and CoIP/MS is available on GitHub (https://github.com/LJ24601) under the project repositories OPP_proteomics, ABPP_rocapal, and coipms, respectively. CRISPRi screen analysis code is available via Zotero (https://doi.org/10.5281/zenodo.18870254).

---

**Figure S1.**
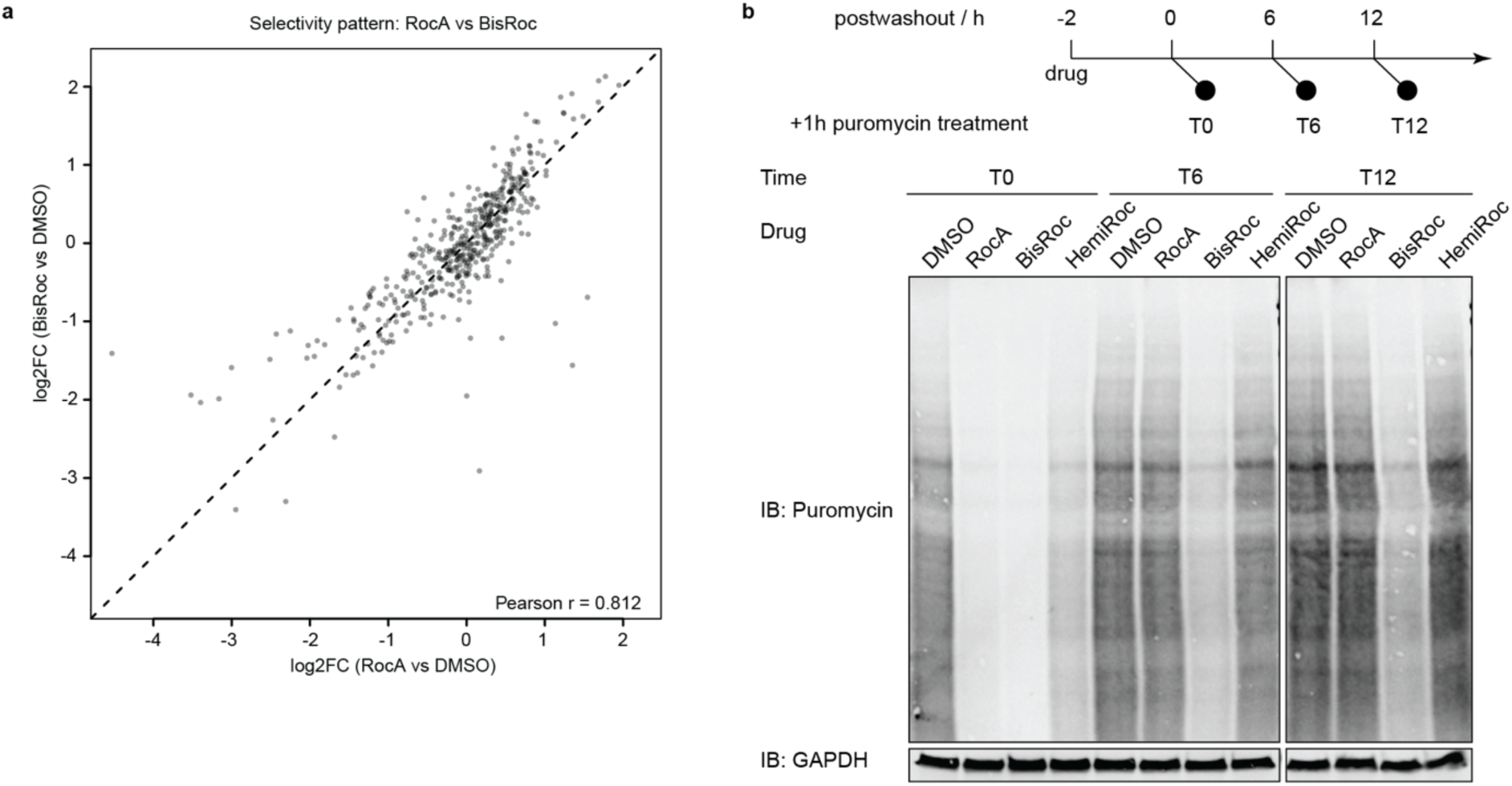
OPP proteomics and washout experiment measuring global translation efficiency using a puromycin incorporation assay. a. Selectivity analysis of nascent protein synthesis by OPP-based quantitative proteomics in K562 cells treated with 100 nM RocA or BisRoc, showing strong concordance in log2 fold-changes relative to DMSO (Pearson r = 0.812). b. K562 cells were pretreated with 1 µM of indicated drugs for 2 h and washed three times with drug-free medium (5 min each), and then incubated for the indicated post-washout times. Puromycin was subsequently added and allowed to incorporate for 1 h before analysis.

**Figure S2.**
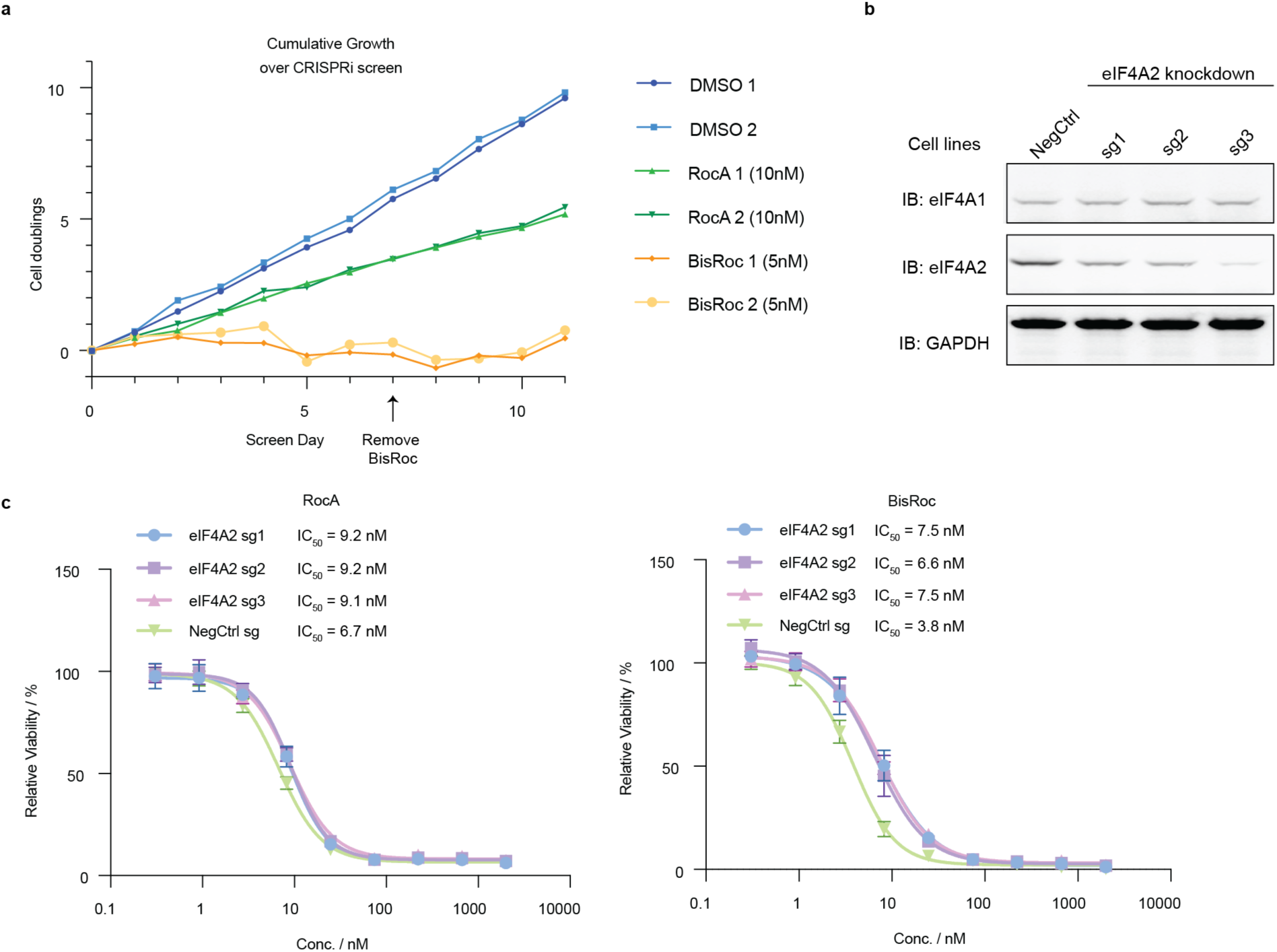
Cell growth during CRISPRi screen and validation of eIF4A2 dependency. a. Cumulative growth over the CRISPRi screen, shown for two independent replicates. b. Western blot detection of eIF4A1 and eIF4A2 levels in the generated stable eIF4A2 knockdown cell lines using sg1-3. c. Validation of eIF4A2 dependency by CellTiter-Glo viability assay performed on eIF4A2 knockdown cell lines (eIF4A2 sg1-3) or negative-control cell line. Data are presented as mean ± s.d. (n = 3).

**Figure S3.**
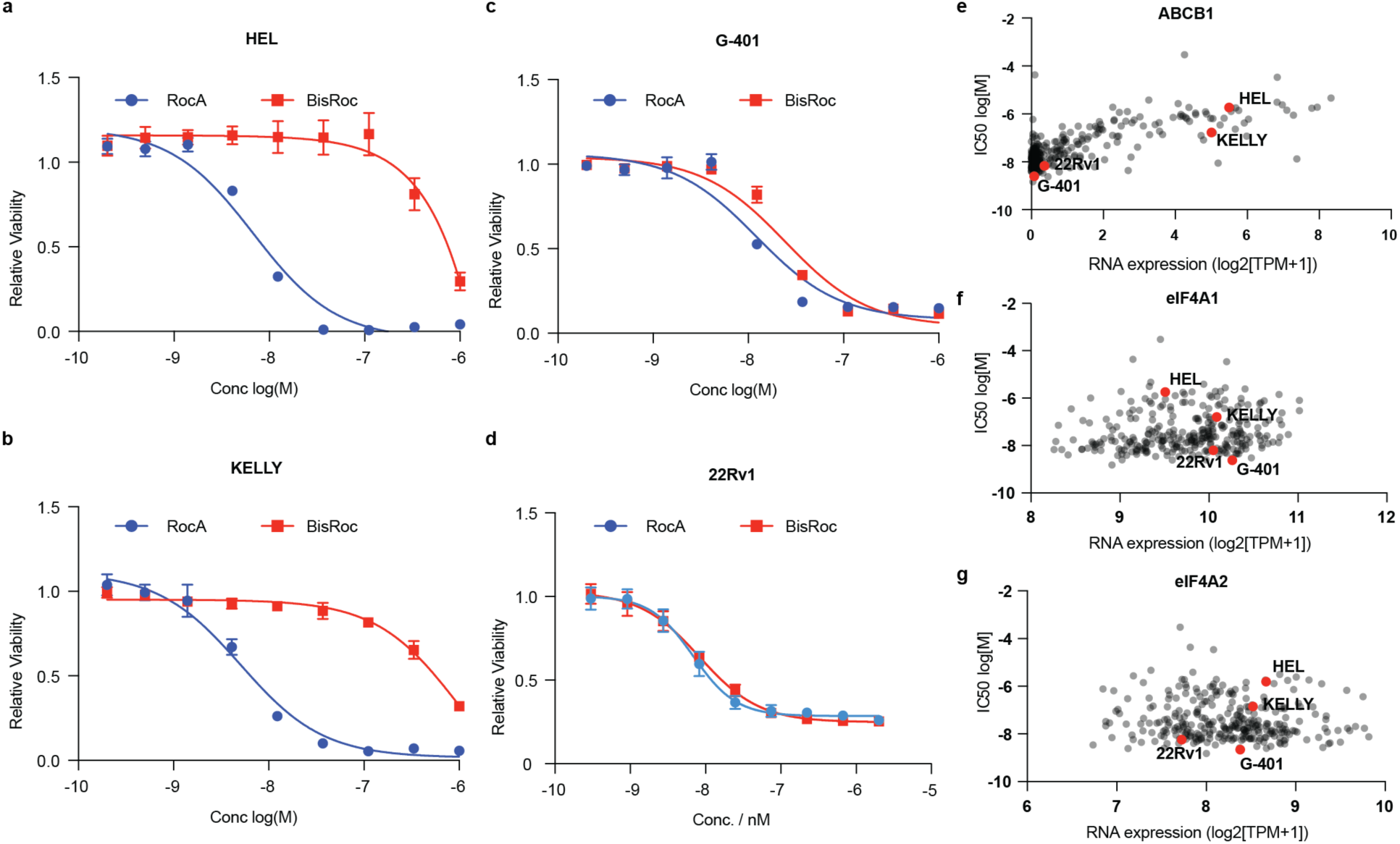
Validation of differential cellular responses to RocA and BisRoc. a–d, Cell viability measured by CellTiter-Glo (CTG) assays. Data are presented as mean ± s.d. (n = 3). e-g, Correlation between BisRoc IC_50_ values and transcript abundance of ABCB1, eIF4A1, and eIF4A2.

**Figure S4.**
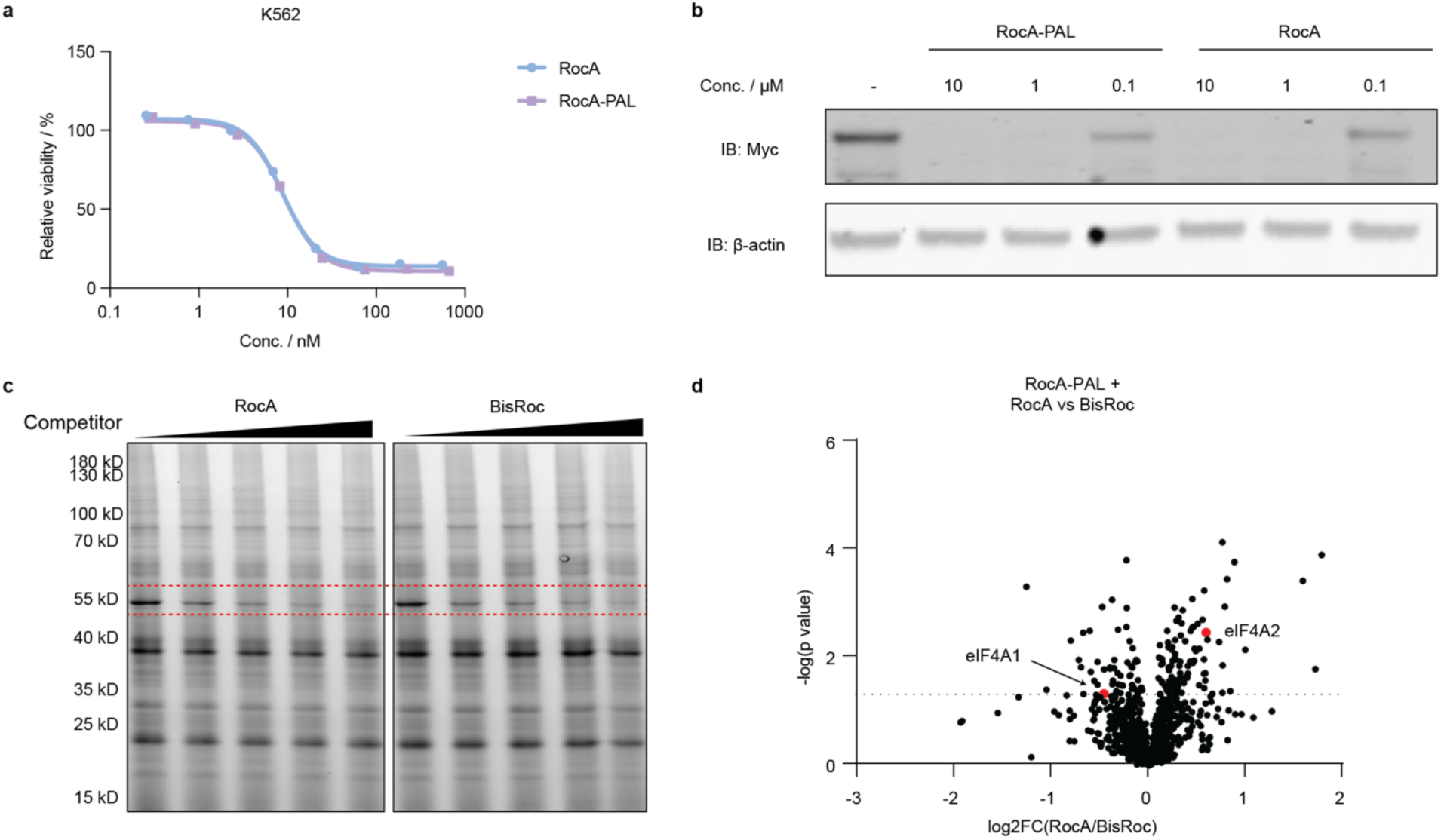
Activity-based proteomic profiling experiment. a, b. RocA-PAL possesses effects comparable to RocA on cell viability and Myc downregulation. Viability data are presented as mean of three independent replicates. c. TAMRA scan of proteome treated with RocA-PAL (5 µM) in combination with increasing concentrations of competitors RocA or BisRoc (0, 5, 10, 20, 50 µM) for 1 h. d. Volcano plot comparing protein enrichment between RocA and BisRoc competition conditions in the presence of RocA-PAL.

**Figure S5.**
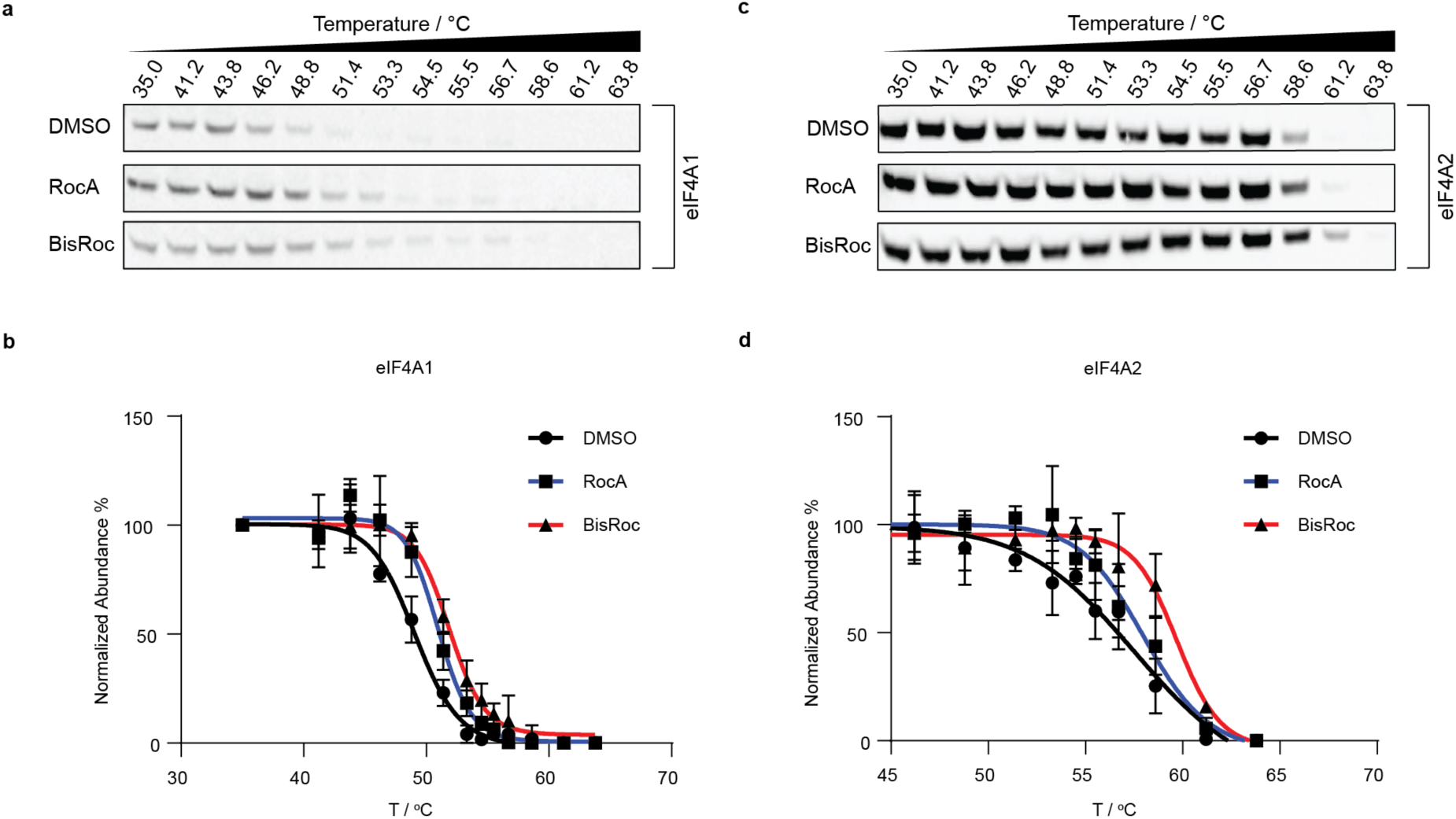
Cellular thermal shift assay (CETSA) for RocA and BisRoc. a, b. Western blot and quantification of CETSA results assessing the target engagement of eIF4A1. Cells were incubated with indicated compounds (10 µM) for 1 h and then heated at the indicated temperature for 3 min. Data are presented as mean ± s.d. (n = 3). c, d. Western blot and quantification of CETSA results assessing the target engagement of eIF4A2. Data are presented as mean ± s.d. (n = 3).

**Figure S6.**
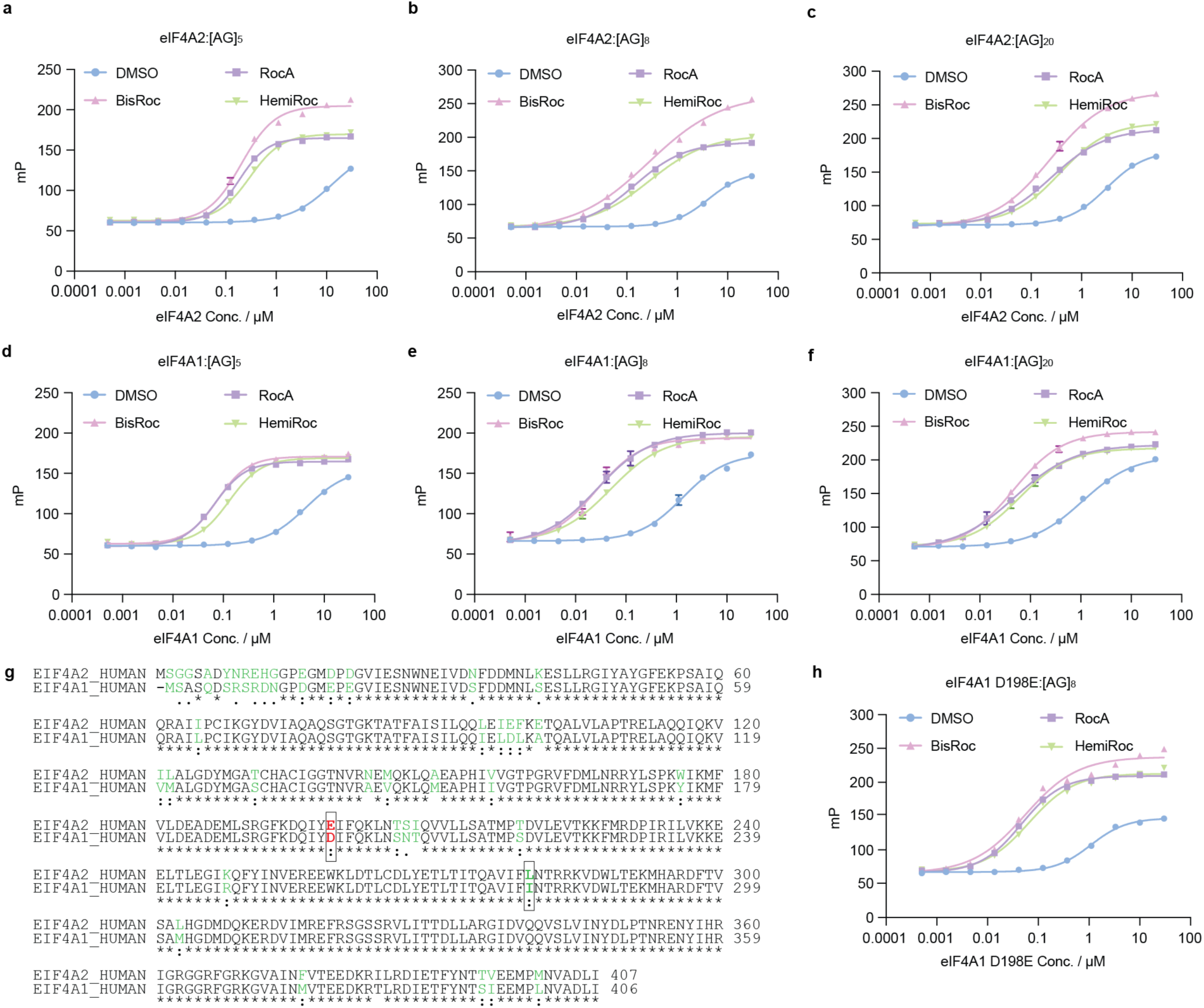
Fluorescent polarization assay and sequence alignment of eIF4A1 and eIF4A2. a-f, Fluorescent polarization assay using eIF4A2 or eIF4A1 at various concentrations and polypurine RNA sequences of varying lengths. Data are presented as mean ± s.d. (n = 3). g. sequence alignment of eIF4A1 and eIF4A2. h. Fluorescent polarization assay using eIF4A1 D198E at various concentrations and polypurine RNA sequence [AG]_8_. Data are presented as mean ± s.d. (n = 3).

**Figure S7.**
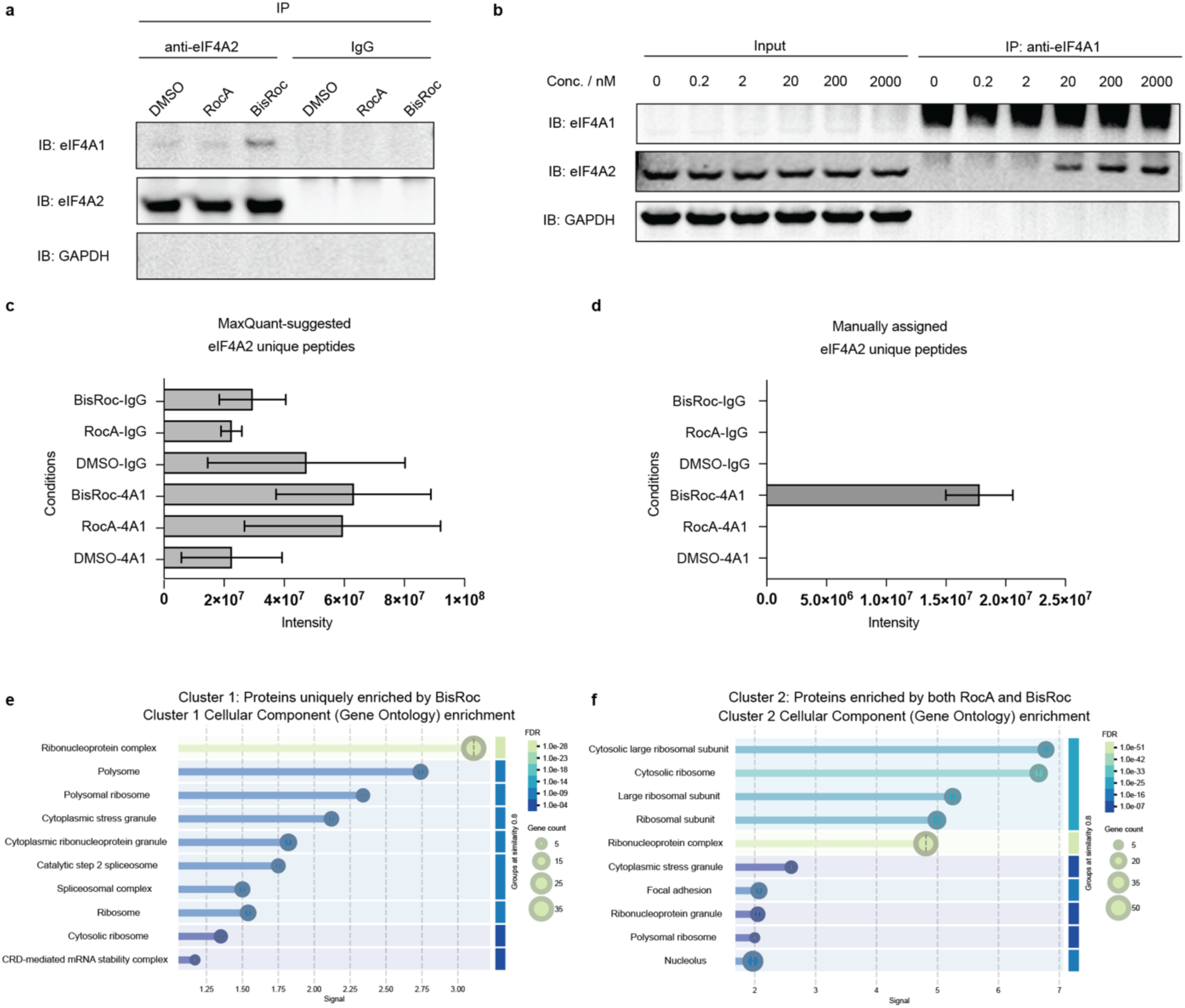
CoIP experiment for BisRoc treatment. a. anti-eIF4A2 co-immunoprecipitates eIF4A1 upon BisRoc treatment. b. Dose-dependent CoIP experiment across a gradient of BisRoc concentrations (2000 nM to 0 nM). An anti-eIF4A1 antibody was used for immunoprecipitation. Data are representative of two independent replicates. c. Intensities of MaxQuant-identified eIF4A2 unique peptides. d. Intensities of manually assigned eIF4A2-unique peptides after excluding four I/L-containing peptides. e. Pathway analysis of the protein cluster 1 uniquely enriched by BisRoc. f. Pathway analysis of the protein cluster 2 commonly enriched by RocA and BisRoc.

**Figure S8.**
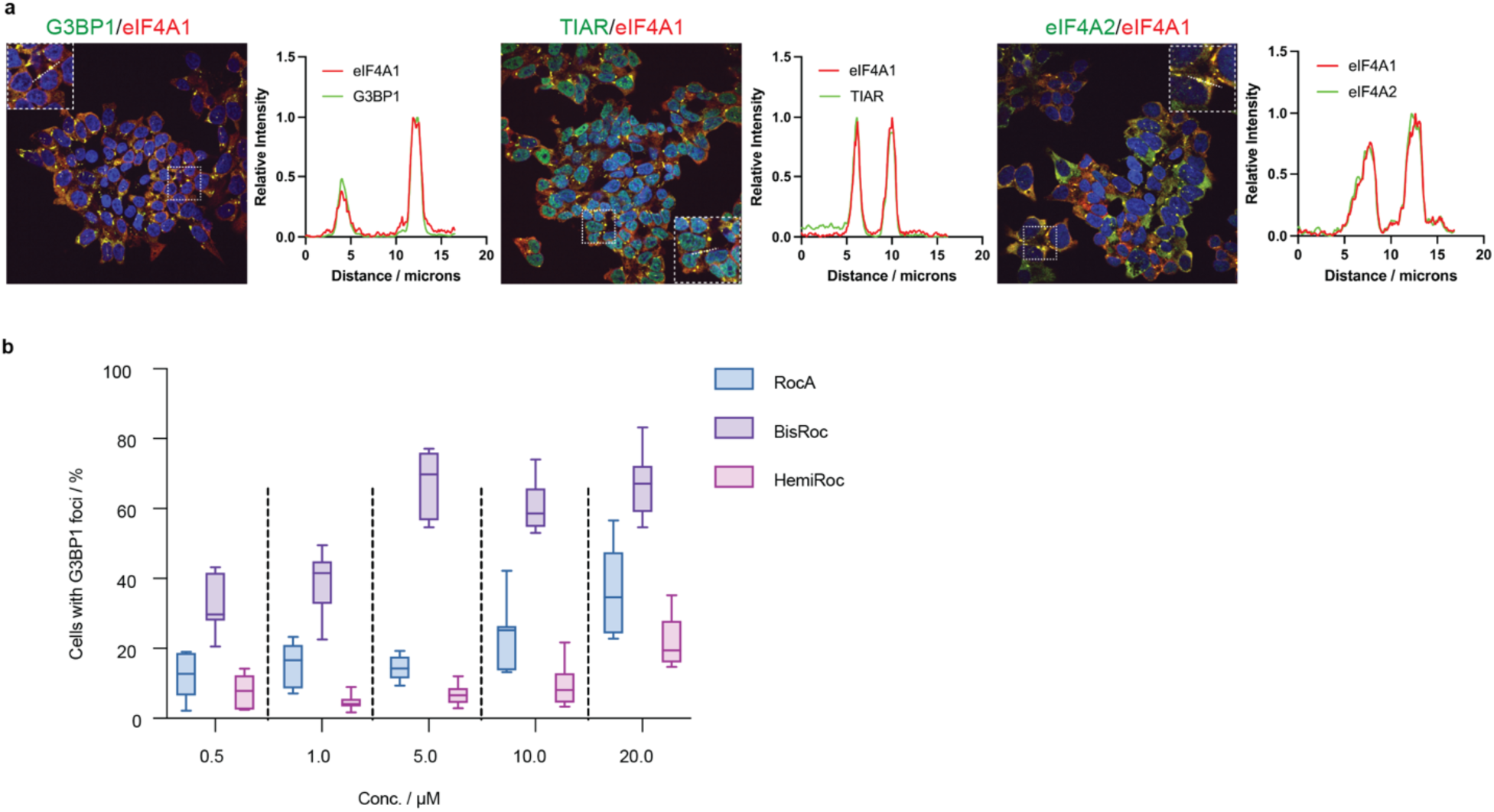
Immunofluorescence microscopy of cells treated with RocA or BisRoc at various concentrations. a. eIF4A1 co-localizes with G3BP1, TIAR, and eIF4A2 under the treatment of 1 µM BisRoc. b. Quantification of G3BP1 foci-positive cells across the range of drug concentrations tested. Data are presented as mean ± s.d. (n = 3).

